# Native architecture of a mitochondrial oxphosome

**DOI:** 10.64898/2026.11.14.766655

**Authors:** Mariia Melnik, Chelsea M. Brown, Siewert J. Marrink, Adrià Nogales Moral, Manon Demulder, Benjamin D. Engel, Florent Waltz

**Affiliations:** Biozentrum, University of Basel, Spitalstrasse 41, CH-4056 Basel, Switzerland; Department of Chemistry, University of Warwick, Coventry, UK; Groningen Biomolecular Sciences and Biotechnology Institute, University of Groningen, Groningen, Netherlands; Gregor Mendel Institute, Austrian Academy of Sciences, Vienna BioCenter, Vienna 1030, Austria

## Abstract

Mitochondrial cristae are the primary bioenergetic units of eukaryotic cells, yet the molecular principles that dictate their morphologies remain largely unknown. Using *in situ* cryo-electron tomography and integrative modeling, we resolve the native organization of the respiratory machinery in the diatom *Phaeodactylum tricornutum*. Unlike previously studied animals and green algae, diatoms possess tubular crista, a conserved membrane architecture found in diverse and globally relevant species. We identify several native supercomplexes assemblies, including an oxphosome comprising the entire respiratory chain. Our results, together with molecular dynamics analyses, reveal that these supercomplexes assemble through a lipid-mediated mechanism. We propose that tubular cristae are shaped by the interplay between membrane-bending ATP synthase oligomers that bud the hemispherical crista tips and supercomplexes that act as molecular roadblocks to constrain ATP synthase propagation, thereby balancing membrane curvature and allowing cristae body extension. Furthermore, we show that the diatom Complex I retains ancestral subunits related to fatty acid synthase. Beyond illuminating diatom bioenergetics, our findings reveal a fundamental principle of mitochondrial morphogenesis where ATP synthase-driven membrane curvature is spatially fine-tuned by respiratory supercomplexes.

---

Mitochondria are essential eukaryotic organelles and the primary sites of molecular respiration, where most of the cell’s adenosine triphosphate (ATP) is regenerated^1–3^. One key feature of their bioenergetic function is the organization of their inner membrane (IM)^4,5^. The folds of the IM, known as cristae, scaffold the five large membrane-embedded enzymes responsible for oxidative phosphorylation^6^. Cristae concentrate respira-tory complexes (complexes I to IV) and ATP synthases (also known as complex V; together hereafter referred to as Complexes (C) I to V) into confined domains, presumably to optimize proton flux and energy conversion^7,8^. Across species and tissues, cristae can adopt strikingly different morphologies^9^. One key determinant of cristae architecture is the supramolecular organization of the respiratory chain itself. Most respiratory complexes associate into higher order supercomplexes, often termed “respirasomes”, whose stoichiometry varies between species, as shown natively^8^ and *ex vivo*^10–12^. Likewise, ATP synthases can dimerize and oligomerize to form rows along the highly curved edges of lamellar cristae, imposing positive curvature and defining their characteristic shapes^13–15^. Together, supercomplex assembly and ATP synthase oligomerization provide a structural basis for sculpting the IM’s three-dimensional topology.

While we recently characterized the native organization of respiratory complexes in green alga^8^, the vast diversity of mitochondrial membrane architecture suggests significant structural variation across eukaryotic lineages. Leveraging environmental data from the TREC expedition^16^ as well as species that were obtained from culture collections, we surveyed species across the protist clades Telonemia, Stramenopila, Alveolata, Rhizaria (TSAR), and Haptista. We combined cryo-focused ion beam (cryo-FIB) milling with *in situ* cryo-electron tomography (cryo-ET) to reveal that these clades share a characteristic tubular cristae architecture (Fig. 1a-d and Supplementary Fig. 1). This morphology sometime serves as a structural hallmark for eukaryote classification^9,17,18^, distinguishing these lineages from the lamellar cristae prevalent in Opisthokonta and Viridiplantae. To investigate the molecular basis of this tubular architecture, we focused on diatoms, secondary endosymbiont algae within the Stramenopile clade^19^. Diatoms are among the most ecologically successful groups of phytoplankton, playing a pivotal role in global carbon fixation and marine food webs^20^. Furthermore, diatoms rely on an intimate energetic coupling where mitochondria actively support chloroplast function, a relationship that is fundamental to their ecological dominance^21^. Despite their ecological and evolutionary importance, little is known about how diatom mitochondria are organized. Here, we use the model diatom *Phaeodactylum tricornutum*^22–24^ to explore the structural principles shaping tubular cristae.

**Figure 1:**
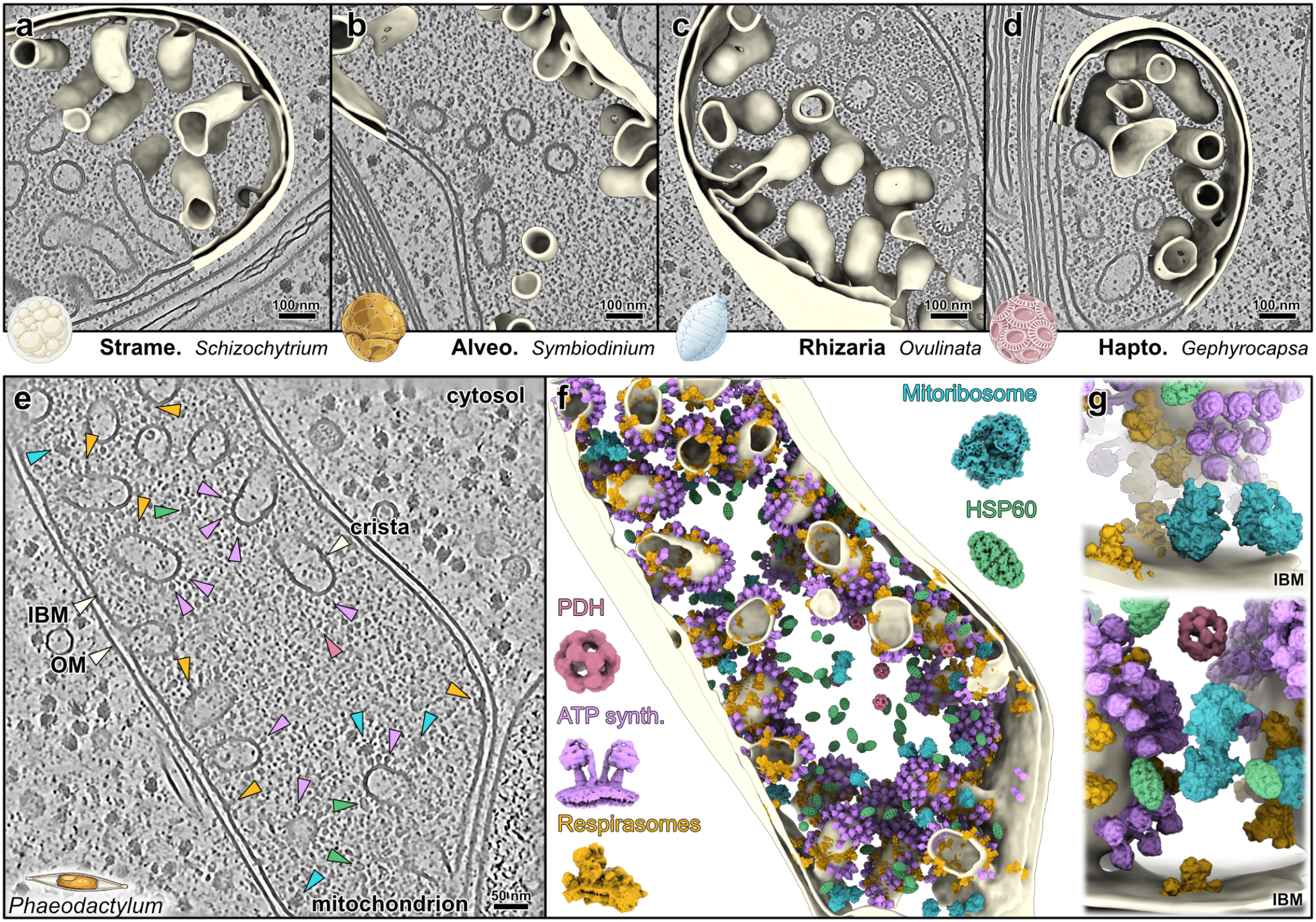
Native molecular architecture of mitochondria with tubular cristae revealed by *in situ* cryo-ET. **a**–**d**, Tomogram slices, all shown at the same scale, and corresponding membrane segmentation of mitochondria across diverse lineages exhibiting tubular cristae architecture: *Schizochytrium sp*. (Stramenopiles) (**a**), *Symbiodinium microadiaticum* (Alveolata) (**b**), *Ovulinata parva* (Rhizaria) (**c**), and *Gephyrocapsa huxleyi* (Haptophyta) (**d**). More examples from different species are shown in (Supplementary Fig. 1). **e**, Tomogram slice depicting a mitochondrion within a native *Phaeodactylum tricornutum* diatom cell (Stramenopiles), showing inner boundary membrane (IBM), outer membrane (OM), and tubular cristae. Colored arrowheads indicate individual macromolecular complexes mapped in **f. f**, 3D segmentation of the tomogram in **e** with subtomogram averages (STA) mapped back into their corresponding spatial coordinates. Beige, mitochondrial membranes; purple, ATP synthase dimers; cyan, mitoribosomes; yellow, respirasomes (some part of oxphosomes); red, PDH; green, HSP60. **g**, Zoomed-in views at the IBM, showing respirasomes found at the IBM and on the cristae, as well as membrane-bound and soluble mitoribosomes and in the PDH and HSP60 in the matrix. Scale bars, 100 nm in **a-d** and 50nm in **e**.

## Diatom mitochondria ultrastructure

Using our *in situ* cryo-ET workflow (see methods), we directly visualized native membrane architecture with molecular resolution inside *P. tricornutum* cells (Fig. 1e-f and Supplementary Fig. 2). In this organism, mitochondria form a continuous network^25,26^, with the distance between the outer membrane (OM) and IM measuring approximately ∼12 nm, which appears to be a general trend across eukaryotes^8^, while the characteristically tubular cristae lumen has an average width of ∼54 nm. This is significantly wider than what is observed in systems with lamellar cristae (∼12-15 nm), such as Chlamydomonas and animals^8,27,28^. Inspection of denoised tomograms allowed direct visualization of several characteristic mitochondrial complexes (Fig. 1e). Leveraging template matching, we accurately localized major soluble complexes within the mitochondrial matrix, including mitochondrial ribosomes (mitoribosomes), the chaperone HSP60, and the Krebs cycle enzyme pyruvate dehydrogenase (PDH) complex (Supplementary Figs. 2 to 4). Detailed analysis of the mitoribosome population revealed distinct functional states, capturing them as free monosomes, hibernating dimers, and membrane-bound complexes (Supplementary Fig. 3). Notably, mitoribosomes were excluded from the cristae regions, like previously observed in green algae^8,29^. However, the presence of free soluble mitoribosomes contrasts with Chlamydomonas and animals, where translation almost exclusively happens at the membrane. This abundance of free mitoribosomes correlates with the *P. tricornutum* mitochondrial genome, which encodes a high proportion of soluble proteins rather than strictly hydrophobic membrane subunits^30^. Within the matrix, PDH complexes were sparsely distributed and exhibited no preferential localization (Fig. 1f). This low copy number may reflect the high catalytic efficiency of the enzyme.

## Tubular cristae accommodate respiratory supercomplexes of variable stoichiometry and membrane curvature

Beyond these soluble complexes, we also identified the major bioenergetic machineries embedded within the inner mitochondrial membrane. By iterative template matching for CI and then I_1_III_2_, we mapped the supramolecular organization of the res-piratory chain within the IM. We resolved the entire chain, CI to CV, revealing that all detected supercomplexes share a con-served I_1_III_2_ core (Fig. 2 and Supplementary Fig. 2). Unlike Chlamydomonas, which contains one main type of respirasome supercomplex, the diatom supercomplexes exhibit striking architectural variability. We classified these into four main supra-molecular assemblies (Fig. 2b and Supplementary Fig. 5): thelargest supercomplex comprising a dimer of the entire respiratory chain 2(I_1_II_1_III_2_IV_1-3_V_2_) and named hereafter oxphosome dimer, a I_1_II_1_III_2_IV_1-3_ respirasome that is similar to the oxphosome but lacks ATP synthases, a dimeric respirasome I_2_III_2_II_1_IV_1-3_ centered around complex III_2_, and a smaller respirasome of I_1_II_1_III_2_IV_1-3_ stoichiometry localized exclusively to the flat inner boundary membrane (IBM). These supercomplexes exhibit different membrane curvature profiles. While the IBM respirasome is localized to a flat membrane geometry, the crista respirasome, oxphosome dimer, and respirasome dimer all occupy cylinder-shaped IM, as evidenced by their membrane topography and principal curvature values (κ_1_ and κ_2_) (Fig. 2c). Furthermore, the dimer respirasome exhibits a slight structural variation that enforces additional curvature (Supplementary Fig. 7f).

**Figure 2:**
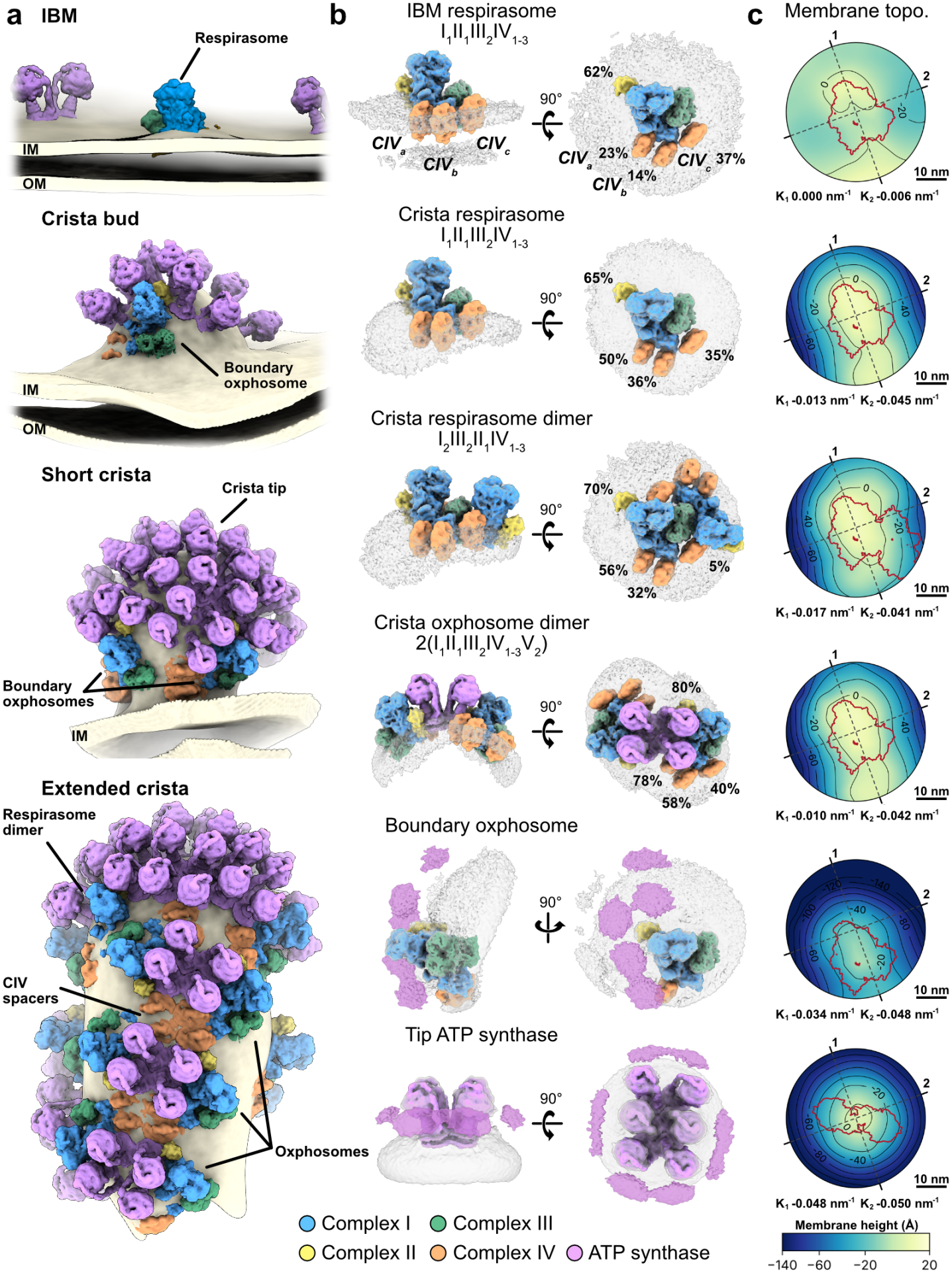
OXPHOS components form supercomplexes of variable stoichiometries in different regions of the inner mitochondrial membrane. **a**, Spatial distribution of oxidative phosphorylation (OXPHOS) complexes across distinct regions of the inner mitochondrial membrane, including the inner boundary membrane (IBM), a putative crista bud, and extended tubular cristae. This may illustrate the progressive stages of crista assembly and shows the partitioning of respiratory complexes and ATP synthase dimers (purple) along varying membrane geometries. Additional examples are shown in Supplementary Fig. 7a-c. **b**, Structural gallery of the main identified respiratory supercomplexes (low passed focused-refined complexes fitted in the supercomplexes STA), presented in side and top views (left and right) with their corresponding membrane environment (transparent gray). Identified assemblies include the IBM respirasome I_1_II_1_III_2_IV_1-3_, crista respirasome I_1_II_1_III_2_IV_1-3_, crista respirasome dimer I_2_III_2_II_1_IV_1-3_, crista oxphosome dimer 2(I_1_II_1_III_2_IV_1-3_V_2_), boundary oxphosome I_1_II_1_III_2_IV_1-3_V_x_, and tip-localized ATP synthase. The occupancy for complexes II and IV within each supercomplex class are indicated in percentages. Individual components are color-coded: complex I (blue), complex II (yellow), complex III (green), complex IV (orange), and ATP synthase (purple). **c**, Membrane topography of each class. Height of the membrane mid-surface (Å) is viewed from the matrix side on a common color scale, same view as in **b**, top view, with all views centered on I_1_III_2_. Contours are marked every 20 Å, and the red outline is the footprint of I_1_III_2_, I_2_III_2,_ or ATP dimer for orientation purpose. κ_1_ and κ_2_ are the curvatures (1/radius) of the mid-surface along axes 1 and 2 (dashed).

We also localized ATP synthases via template matching (Supplementary Figs. 2 and 6). In contrast to *C. reinhardtii*, where respiratory complexes and ATP synthases occupy different membrane domains (flat and curved respectively), the tubular cristae of *P. tricornutum* accommodate both assemblies in close spatial proximity, integrated within the same supercomplex— the oxphosome—and as separate standalone assemblies. Within the oxphosome dimers occupying the cylindrical cristae body regions, ATP synthases form tetrameric units (arranged as a dimer of dimers) (Fig. 2b), which were often observed organized into higher-order rows aligned longitudinally along the cristae body (Fig. 2a). Mapping nearest neighbors for all respirasome positions confirmed this pattern, revealing a regular arrangement that directly matches the oxphosome rows (Supplementary Fig. 7a and d). In contrast, the crista tips are exclusively occupied by extended oligomers of ATP synthases that generate the high membrane curvature of these hemispherical tip regions (Fig. 2 and Supplementary Fig. 6). At the boundary where the cristae body transitions into the ATP synthase-en-riched cristae tip, we found respirasomes that were always oriented the same way, with CII and the CI matrix arm facing toward the cristae tip and the CI membrane arm pointing towards the cristae body. While systematically found at this position, the association between respirasome and ATP synthases is weak and prevented determining a clear stoichiometry for ATP synthase relative to the respirasome core. We list it as I_1_II_1_III_2_IV_1-3_V_x_. Because these respirasomes and their attached ATP synthases delineate the boundary between crista body and tip, we name them boundary oxphosomes (Fig. 2 and Supplementary Fig. 5).

All detected supercomplexes exhibited substantial stoichiometric and structural variability, reflected in the diverse states resolved by 3D classification. While CI-CIII association remained stable, CII and CIV exhibited flexible association with CI across all super-complex populations, posing challenges for focused refinement of CII and CIV (Supplementary Fig. 8). We identified three distinct CIV binding sites (CIV_a_CIV_b_and CIV_c_), characterized by varying levels of stability depending on the membrane curvature. CIV_”_occupies a proximal position relative to the CI matrix arm, positioned close to the carbonic anhydrase module, analogous to the ‘M’ site in *Chlamydomonas*^8^ (Supplementary Fig. 9a). In oxphosome dimers, CIV_a_is situated next to the integrated ATP synthase te-tramers. CIV_”_anchors adjacent to CIV_”_, with the same orientation, exhibiting variable occupancy and potential spatial competition with other complexes (Fig. 2b and Supplementary Fig. 8). Notably, both CIV_a_and CIV_b_maintain higher occupancy exclusively within cristae-localized supercomplexes, dropping to residual levels (below 23%) in the flat IBM class. Conversely, CIV_c_binds at the distal tip of the CI membrane arm across all super-complexes, as in mammalian counterparts^11,31^ (Supplementary Fig. 9a), and exhibits similar occupancy in super-complexes localized to both flat and curved membrane regions. An exception is the respirasome dimer, where the second CI probably negatively impacts its occupancy (down to 5%). Due to pronounced positional heterogeneity relative to CI, CIV_”_was the least well resolved. The most stable CIV association was observed for CIV_a_and CIV_b_within the oxphosome dimers, where a highly organized “doublet” of CIV_a-b_(up to 78% occupancy) appears to act as a structural spacer for ATP synthase tetramers, promoting regularity of oxphosome rows (Fig. 2a, Extended crista and Supplementary Fig. 7a). Notably, none of CIV_a-c_appear to form a defined protein-protein interaction with CIII_2_, while only CIV_”_directly contacts CI (Supplementary Fig. 9b). CII associates with the back of the CI proximal membrane arm, exhibiting a sub-stoichiometric occupancy of approximately ∼62-80% across the total supercomplex population, which is higher compared to CIV_”_(Fig. 2b and Supplementary Fig. 8). Like CIV, CII associates flexibly with CI, but predominantly adopts two discrete orientational states relative to it (Supplementary Fig. 8). This structural heterogeneity may reflect local molecular crowding; the specific orientation of CII is likely restricted by steric constraints and the presence or absence of adjacent complexes within the densely packed membrane environment.

## Integrative approach elucidates the native structure of the respiratory chain

Using focused refinement on each complex, we managed to reach resolutions of 4.5 Å on CI, 10 Å on CII, 6.4 Å on CIII_2_, 10 Å on CIV, and 9 Å on CV, including the four main ATP synthase catalytic states (Supplementary Figs. 5, 6 and 8). To assemble atomic models, we initially attempted isolating mitochondria and respiratory supercomplexes for single-particle analysis. However, mitochondria purification was hampered by their tight physical association with the secondary chloroplast^21,26,32^, and the supercomplexes dissociated during biochemical isolation (Supplementary Fig. 10). Conse-quently, *in situ* cryo-ET proved the only viable technique to resolve the intact architecture of the supercomplexes. Hence, we opted for an integrative approach. We identified the different protein subunits of the complexes based on mass-spectrom-etry (Supplementary Fig. 10), combined with sequence and structural homology searches via BLAST^33^ and FoldSeek^34^ and then generated the initial models using AlphaFold3 multimer^35^, subsequently refined in the subtomogram averaged maps (Supplementary Figs. 11-12). This allowed us to build near-complete models of entire diatom respiratory chain, with CI composed of 51 proteins, CII of 4 proteins, CIII_2_ of 10 proteins,CIV of 11 proteins and ATP synthase of 14 proteins (Fig. 3, Supplementary Figs. 11-12 and Supplementary Table 2). In general, these complexes are particularly conserved with the rest of eukaryotes, notably CII and CIII_2_, which are nearly identical in terms of structure and number of protein subunits to plants, yeast, and animal complexes^36–39^.

**Figure 3:**
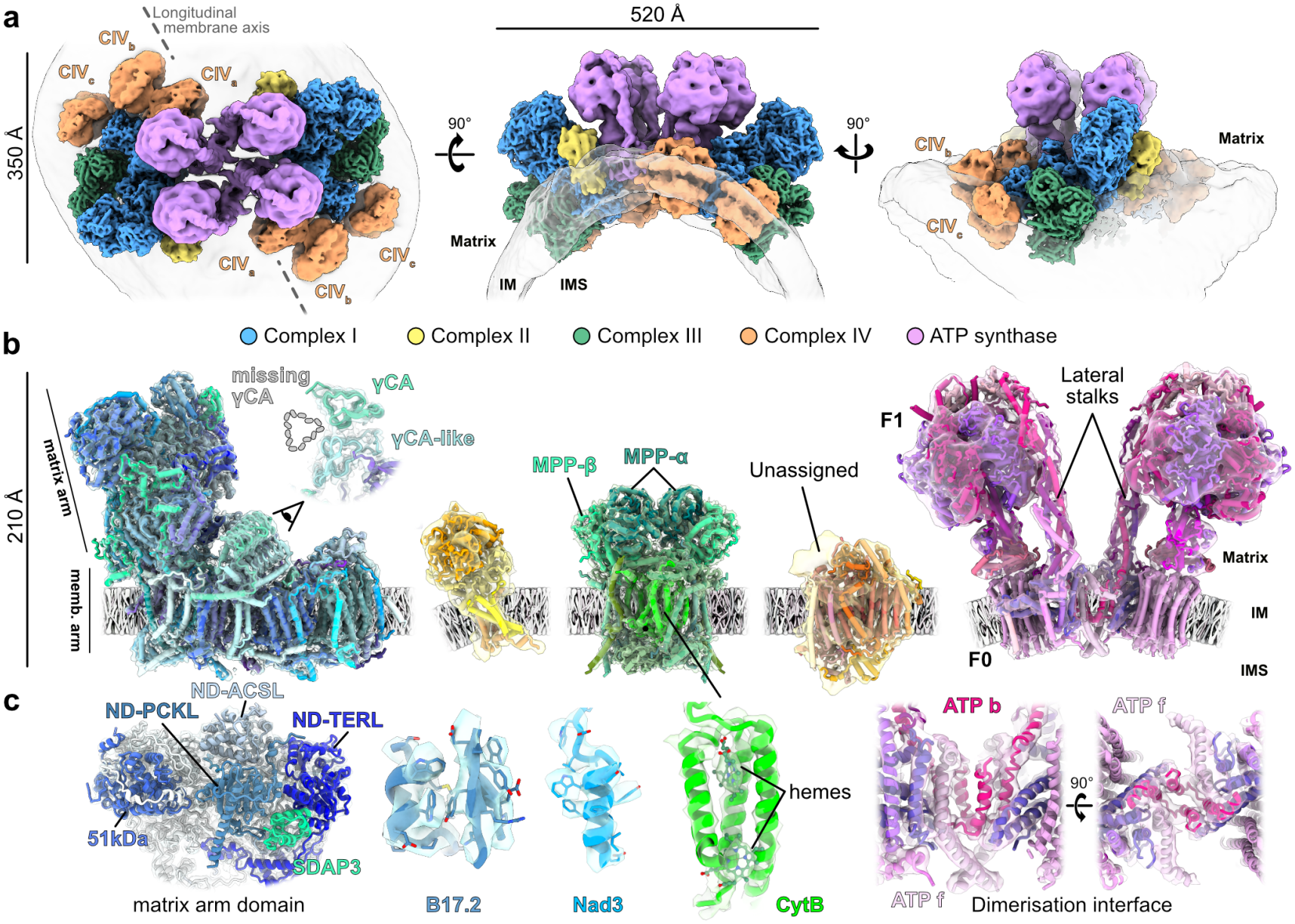
Native structure of the diatom oxphosome resolved by an integrative approach. **a**, Subtomogram averaging (STA) maps of the *P. tricornutum* oxphosome dimer, shown as a composite of focused refined respiratory complexes. From left to right: view from the matrix side, side view shown from along the longitudinal membrane axis, and a 90° rotated view. **b**, Integrative atomic models of the respiratory chain complexes fitted into their respective STA density maps, shown in transparency. Complexes are displayed to scale in cartoon representation (from left to right): complex I (blue shades), complex II (yellow shades), complex III (green shades), complex IV (orange shades), and ATP synthase (purple shades). Complete subunit compositions are detailed in Supplementary Figs. 11, 12 and Table 2. **b**, Close-up views highlight structural details, including specialized auxiliary sub-units within the complex I matrix arm domain, atomic model fits for specific protein components and cofactors such as hemes in complex III, and the structural interface mediating ATP synthase dimerization. With IM inner membrane and IMS inter-membrane space.

The ATP synthase dimer adopts a Type III-like architecture, sharing the overall dimer orientation and lateral stalk positioning characteristic of Alveolata, but with a ∼30° angle between the monomers. However, unlike the massive, highly divergent complexes of *Tetrahymena*^40^ and *Toxoplasma*^41^, dimerization is mediated primarily by subunits ATP b and ATP f, with the latter likely playing a role in higher-order oligomerization (Fig. 3c, Supplementary Fig. 13). This represents a minimal baseline variant of the Type III architecture within the SAR supergroup. CI presents some of the most characteristic features among these complexes. Its 75-kDa core subunit is fragmented, and its inactive carbonic anhydrase (CA) domain is dimeric rather than the trimeric form found in other species (Fig. 3c, Supplementary Fig. 11). A defining feature of the diatom CI is the large additional domain anchored to the matrix arm tip, which is composed of four subunits: ND-TERL, SDAP3, ND-PCKL, and ND-ACSL (Fig. 3c). Intriguingly, these additional subunits are orthologous to those identified at the identical structural position in *Euglena gracilis*^42^, with one subunit (B7G0K8) also conserved in *Tetrahymena*^10,43^ (Fig. 3b and Supplementary Fig. 14).

ND-TERL, SDAP3, and ND-ACSL exhibit sequence homology to proteins involved in fatty acid metabolism, while ND-PCKL connects phosphoenolpyruvate and oxaloacetate metabolism. They are, however, not active (Supplementary Fig. 14). Given that diatoms and euglenids are separated by at least 1.5 billion years of evolution^44,45^, the independent acquisition of four identical subunits at the same exact position on CI seems improbable. Instead, we propose that these fatty acid-related subunits represent ancestral eukaryotic components of CI that were subsequently lost in other eukaryote lineages, such as Opisthokonta and Viridiplantae. This evolutionary trajectory mirrors that of the CA domain, which is now recognized as a near-ubiquitous feature of mitochondrial CI, with the exception of the Opisthokonts^38,46,47^.

### Molecular dynamics analysis implicates cardiolipin in oxphosome assembly

One striking feature of the diatom supercomplexes is that, un-like those characterized in other organisms^8,10,11^, their supramolecular assemblies exhibit few protein-protein contacts or, in some cases, none at all (Fig 3a and 4a). This structural plasticity is particularly pronounced for CII and CIV, with their strong positional and occupancy heterogeneity (Supplementary Fig. 9). Furthermore, we identified no direct protein-protein contacts tethering ATP synthase to CI, and similarly for CIV_b_and CIV_a_(Fig. 4a and Supplementary Fig. 9). This is in line with the “minimalist” nature of the individual diatom respiratory complexes, which are almost entirely restricted to the core catalytic subunits plus core mitochondrial subunits and lack extensive accessory subunits found in other lineages.

**Figure 4:**
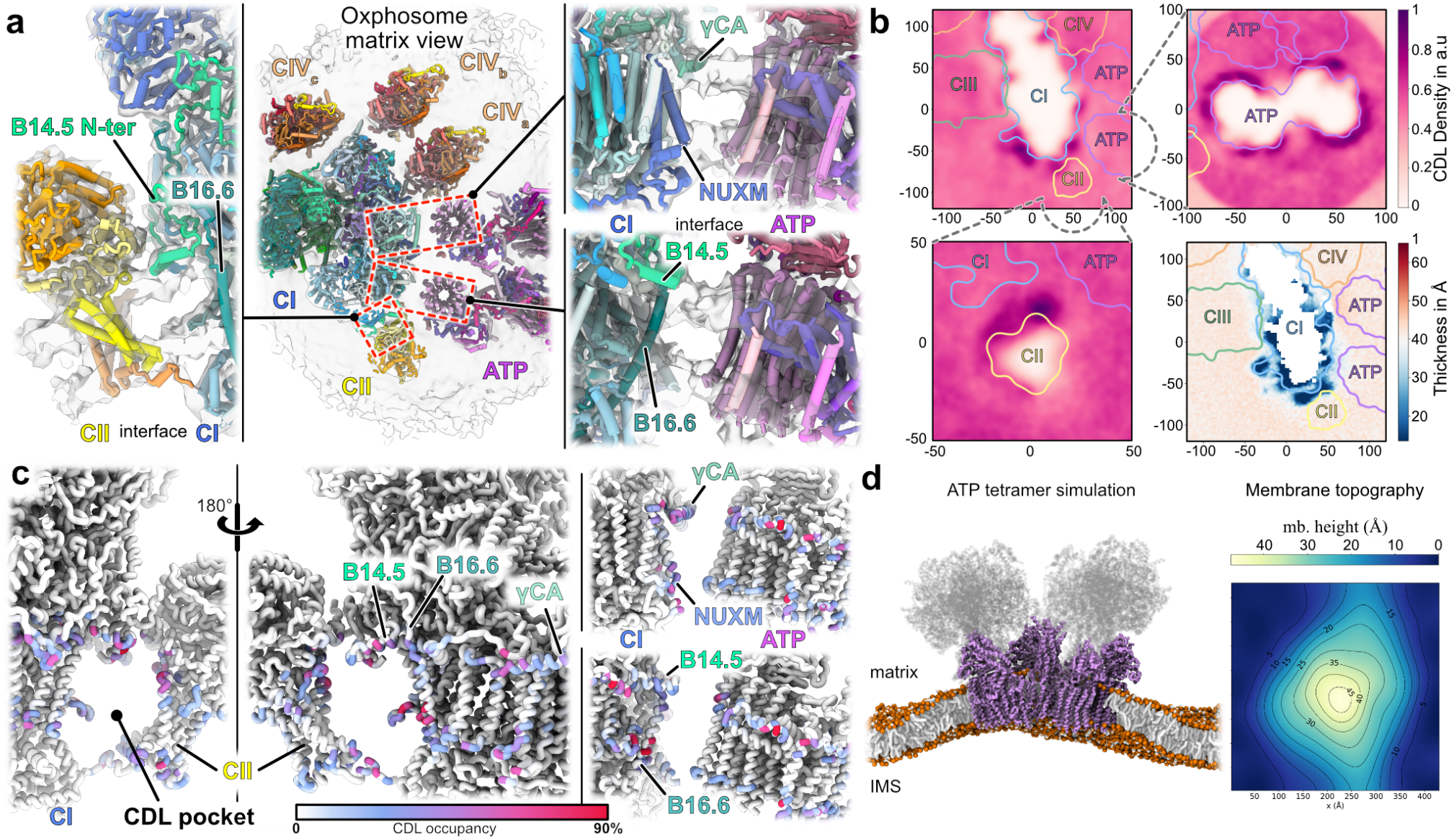
Respiratory supercomplex formation involves membrane thinning and cardiolipins. **a**, Molecular model of the oxphosome viewed from the matrix side, showing the gap between the complexes, notably between CI and the ATP synthase dimers, as well as CIV_b_ and CIV_c_. Close-up panels highlight the CI - CII interface mediated by the B14.5 N-terminus, and the absence of protein-protein interaction at the CI and ATP synthase interface; for each dimer, this is bridged by a 25–30 Å membrane gap. **b**, Molecular dynamics simulations of respiratory complexes, showing cardiolipin (CDL) densities around CI, CII, and ATP synthase, as well as membrane thinning around CI (bottom right panel). Outlines of the different complex positions are shown. All complexes are shown in Supplementary Fig. 15. **c**, Projection of CDL binding sites and occupancy onto the atomic models of CI, CII, and ATP synthase, shown in similar views as in **a** with key interaction pockets highlighted as well as the closest CI subunits to each ATP synthase dimer. **d**, Structural simulation and corresponding membrane topography map showing local membrane bending induced by an ATP synthase tetramer (F1 region, shown in gray, removed for computation efficiency).

To investigate how these complexes are held together, we used molecular dynamics (MD) to study protein complex-lipid interactions. In order to study lipid association with the proteins over a long time scale, the coarse grained (CG) Martini 3 force field^48^ was used. After simulating each complex separately in a bilayer resembling the inner mitochondrial membrane^10^, we measured the density of each lipid species in the surrounding environment. While phosphatidylethanolamine (PE) and phosphatidyl-choline (PC) did not show any enrichment at the protein surface, cardiolipin (CDL) was recruited at distinct sites for all respiratory complexes (Fig. 4b and Supplementary Fig. 15), in line with previous simulation results^49–51^. These binding sites are occupied by CDL in up to 90% of the simulation time (Fig. 4c) and correlate well with the contact sites between these com- plexes, notably CI, CII, and ATP synthase (Figs. 4a and c). In addition to lipid recruitment, membrane perturbation was measured surrounding each of the individual complexes. Substantial membrane thinning was observed around most complexes (Fig. 4b and Supplementary Fig. 15) and could help drive both complex association and membrane curvature. The combination of CDL recruitment and membrane thinning was particularly pronounced around CI, at positions correlated with the location other components of the supercomplexes, notably with CII and ATP synthase. This could explain the lack of direct protein-protein contacts and instead indicate lipid-mediated complex formation.

## Discussion

Here, we investigate the native molecular basis of tubular cristae architecture in the diatom *P. tricornutum*. We describe a *Native architecture of a mitochondrial oxphosome* large assembly composed of the entire respiratory chain, the oxphosome, as well as multiple other flexible supercomplexes of variable stoichiometry, residing on different regions of the mitochondrial IM, whether on the curved cristae membranes or on the flat boundary membrane. The ability to resolve diverse stoichiometries and flexible associations between complexes, while mapping these assemblies to distinct regions of the IM, highlights the power of *in situ* cryo-ET to quantify native protein organization within the native architecture of cellular membranes.

Taken together, our results indicate that individual diatom respiratory complexes are remarkably streamlined, consisting almost exclusively of conserved core subunits and mitochondria-specific proteins found across eukaryotes, with few line-age-specific additions (Fig. 3). Curiously, these complexes assemble into one of the largest known respiratory supercomplexes, the oxphosome (Fig. 2). It is possible that the streamlined nature of the diatom complexes enables the high-degree of supercomplex modularity we observe. Unlike the specialized protein scaffolds in the ciliate Tetrahymena^10,52^ that comprise tens of lineage-specific subunits and protein extensions dedicated to stabilizing the assembly and inducing membrane curvature, the diatom supercomplexes appears to dispense with such structural “glue” (Fig. 4). Instead, the association is inherently flexible and primarily lipid-mediated, as evidenced by the significant range of motion observed for CII and CIV relative to the I_1_III_2_ core.

Using MD, we find that CDL, the signature lipid of the mito-chondrial inner membrane^53^, are enriched at the interface between complexes, with CI serving as a central connecting hub within the supercomplex. In addition, we see that local membrane thinning occurs together with this CDL enrichment. We propose that local protein concentration, coupled with specific lipid recruitment and induced membrane thinning, drives the association of respiratory complexes without requiring direct, high-affinity protein–protein contacts. This is in line with the proposed model for mammalian supercomplex formation^11,54^. Additionally, we see that lateral CIV_”_and CIV_”_are present at higher occupancy in supercomplexes localized on curved membranes compared to those on flat membranes (Fig. 2b-c). This suggests that curved membrane topology may favor CIV incor-poration into the supercomplexes rather than the supercomplexes driving membrane curvature. Moreover, only ATP synthase tetramers (Fig. 4d) and higher-order oligomers appear to be able to substantially curve the membrane. Our *in situ* maps of native mitochondria further reveal that respiratory complexes the protons pumps, ATP synthases – the proton sinks are inter-mixed and often closely associated within the cristae. This in-dicates that strict lateral heterogeneity is not required for ATP generation. Nonetheless, the crista tips are exclusively decorated with ATP synthase, which is likely required to impart their high-curvature hemispherical shape. Consistent with previous studies, we propose that ATP synthase oligomers are the main driver of membrane curvature^13,14^, both in this diatom system and generally in mitochondria. However, this needs to be regulated to prevent suboptimal membrane topology, e.g. membrane tubes fully covered by ATP synthases (exemplified in the parasite Trypanosoma^55^) or ATP synthase-coated vesicles budding off the IM. In line with this idea, we observe supercomplexes with consistent orientation at the boundary between the ATP synthase-covered highly curved tip and the cylindrical body, acting as molecular “roadblocks” that limit the extent of ATP synthase oligomerization. We propose that association with respiratory complexes may regulate the oligomerization of ATP synthase and, consequently, membrane architecture. The diameter of the cristae is naturally set by the hemispherical tip specified by ATP synthases. Along the cristae body below the tip, the ATP synthase tetramers within the oxphosome rows promote curvature but are balanced by the associated respiratory complexes to produce cylindrical membrane curvature.

Based on these findings, we propose a hypothetical stepwise model for tubular crista morphogenesis (Fig. 5): 1) Crista bio-genesis initiates through local ATP synthase oligomerization, which drives membrane curvature and budding from the inner boundary membrane (IBM). 2) To prevent unconstrained assembly, respiratory supercomplexes accumulate at the boundary zone, acting as molecular roadblocks that arrest ATP synthase oligomerization and define the hemispherical cap of the crista tip. 3) Crista elongation subsequently proceeds via continuous insertion of additional respirasome and oxphosomes supercomplexes, extending the tubular crista body longitudinally while enforcing cylindrical membrane curvature. 4) Finally, at the crista base, MICOS complexes, though unresolved in our study, presumably coordinate membrane constriction to establish and stabilize the crista junction^56,57^. Beyond diatoms and tubular cristae, these findings point to a fundamental principle of mitochondrial architecture. While ATP synthases drive membrane curvature, their self-assembly must be balanced to meet architectural constraints, with respiratory supercomplexes serving to fine tune crista morphology.

**Figure 5:**
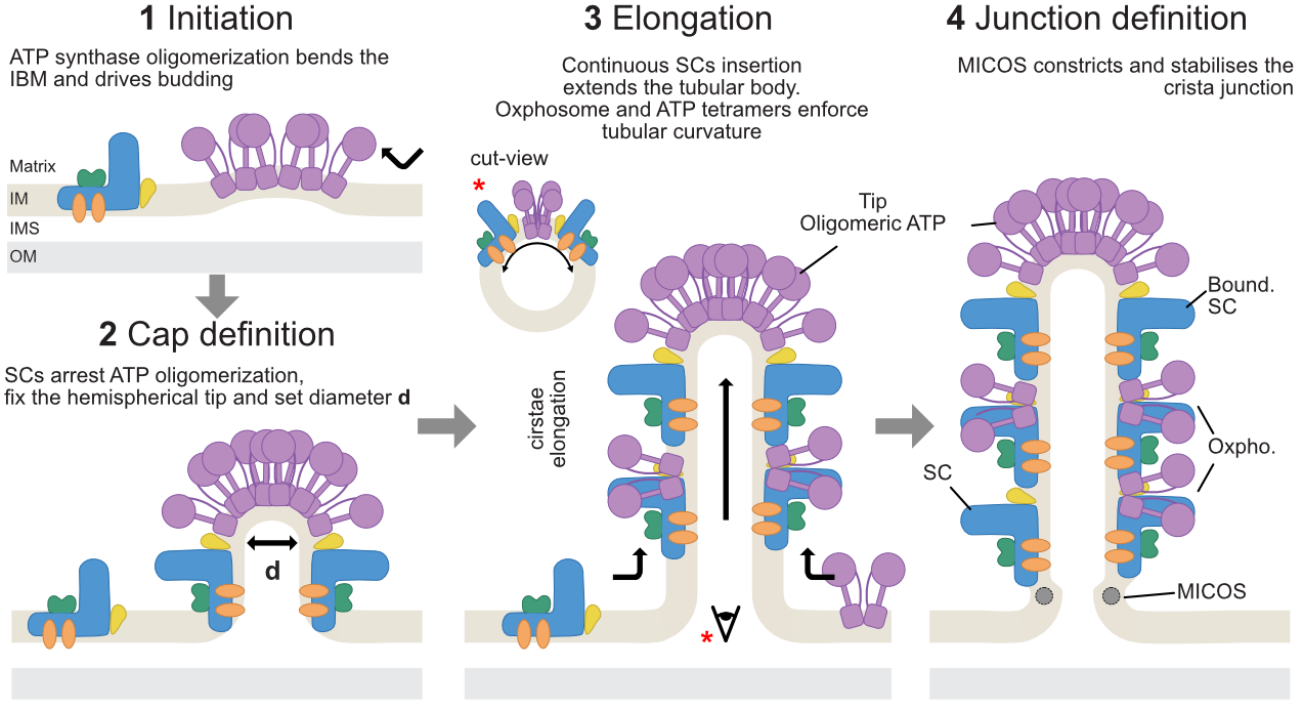
Proposed hypothetical model of tubular cristae formation. Speculative sequential pathway (**1–4**) illustrating tubular crista morphogenesis. Cross-section view along the tubule axis is shown in step **3** (indicated by the red asterisk and eye symbol). Complex color coding is consistent with Fig. 2: complex I, blue; complex II, yellow; complex III, green; complex IV, orange; ATP synthase (‘ATP’), purple. Bound. SC, boundary supercomplex; *d*, tubule diameter; IBM, inner boundary membrane; IM, inner membrane; IMS, intermembrane space; MICOS, mitochondrial contact site and cristae organizing system; OM, outer membrane; Oxpho., oxphosome.

## Data availability

Subtomogram averages of *P. tricornutum* mitochondrial complexes have been deposited in the Electron Microscopy Data Bank (EMDB), and atomic models in the Protein Data Bank (PDB), under the following accession codes. **Focused respiratory complexes:** focused complex I, EMD-100471 (PDB 35FC); focused complex II, EMD-100473 (PDB 35FE); focused complex III, EMD-100472 (PDB 35FD); focused complex IV, EMD-100474 (PDB 35FF). **Respiratory supercomplexes:** IBM respirasome, EMD-100477; crista respirasome, EMD-100478; dimer respirasome, EMD-100700 (PDB 35NO); oxphosome, EMD-100479; composite oxphosome, EMD-100623 (PDB 35KH). **ATP synthase:** unfocused dimer, EMD-100464; composite map, EMD-100483 (PDB 35FH); C2 stalk focus, EMD-100465; State 1, EMD-100466; State 2, EMD-100467; State 3a, EMD-100468; State 3b, EMD-100469; tip class, EMD-100480; oxphosome class, EMD-100475; non-oxphosome, non-tip class, EMD-100476. **Mitoribosome:** consensus reconstruction, EMD-100293; focused LSU, EMD-100294; focused SSU, EMD-100295; membrane-bound class, EMD-100297; hibernating dimer class, EMD-100296. **Other complexes:** HSP60, EMD-100298; pyruvate dehydrogenase (PDH), EMD-100299. **Tomograms:** tomograms shown in Figure 1 are available in the EMDB under accession codes EMD-100284 (*P. tricornutum*), EMD-100285 (*G. huxleyi*), EMD-100286 (*Schizochytrium sp*.), EMD-100287 (*O. parva*) and EMD-100288 (*S. microadriaticum*).

Raw cryo-ET data for *P. tricornutum* are available in the Electron Microscopy Public Image Archive (EMPIAR) under accession code EMPIAR-XXXXX.

## Acknowledgements

We thank C. F. Rodriguez and M. Hunkeler from the University of Basel BioEM facility for their help operating the electron microscopes. We thank sciCORE (https://scicore.unibas.ch/) scientific computing center at University of Basel. We thank D. Ritz from the University of Basel Proteomics Core facility for their assistance with sample preparation and data analysis. We thank E. Dumanoglu and M. Pokutta of the T. Maier group for providing us with the Schizochytrium cells. We thank the Traversing European Coastlines (TREC) Consortium, the TREC core partners of EMBL, the Tara Ocean Foundation, the Tara Europa team and the European Marine Biological Resource Centre (EMBRC) for their commitment to making the TREC-Tara Europa Expedition possible. We thank the Mobile Laboratories Facility of the European Molecular Biology Laboratory for their support. We thank all the members of the Biozentrum Cell Architecture Lab for their valuable input and discussions. We would like to thank SURF for access to Snellius computing resources.

## Funding

M.M. and F.W. and were supported by the Swiss National Science Foundation an SNSF Ambizione grant (project 216094). A.N.M., M.D., and B.D.E. were supported by ERC consolidator grant “cryOcean” (fulfilled by the Swiss State Secretariat for Education, Research and Innovation, M822.00045). C.M.B and S.J.M was supported by European Research Council (ERC) with the Advanced grant 101053661 “COMP-O-CELL”.

## Author contributions

M.M. performed biochemical purifications, cryo-ET data processing, and initial writing of the manuscript. M.M. and F.W. performed cryo-FIB milling, collected cryo-ET data, and interpreted results. A.M.N. and M.D. helped with cryo-ET of environmental samples and culture collections. C.M.B. and S.J.M. performed MD analyses. F.W. and B.D.E. conceived and funded the project. F. W. wrote the manuscript with input from all the authors.

## Competing interests

The authors declare no financial interests/personal relationships that may be considered as potential competing interests.

## Materials and Methods

### Phaeodactylum culture conditions

The *P. tricornutum* Pt1 strain (CCAP 1055/3; Scottish Marine Institute, UK) was kindly provided by S. Flori. Cells were cultured in Enriched Seawater Artificial Water (ESAW) medium^59^ as previously described by Flori et al.^26^. Briefly, cultures were grown and maintained in 100 mL flasks containing a maximum of 40 mL of medium. Cultivation was performed in a growth cabinet (Binder KT170) at 19 °C under a 12-h light/12-h dark photoperiod with a light intensity of 20 μmol photon m^”2^ s^”1^ and continuous shaking at 100 rpm, aerated with ambient air. For large-scale growth, cultures were iteratively scaled up from 100 mL flasks to 2 L flasks (final volume, 1 L of culture).

### In situ cryo-ET

#### Grid preparation

4 μl of cell suspension were applied onto a Quantifoil R2/1 300-mesh holey carbon grid that were glow-discharged at 5W for 20 sec in a Gatan glow discharger. The sample was plunge frozen in liquid ethane with a Leica GP2 system (temperature = 20 °C, humidity 100%) with a wait time of 5 sec before blotting for 4 sec.

#### FIB milling

FIB milling was performed as described in^60^. Briefly, grids clipped in Autogrids (Thermo Fisher Scientific) were loaded into an Aquilos 2 microscope (Thermo Fisher Scientific) and a layer of organometallic platinum was applied using the gas injection system. For milling and imaging, the gallium ion beam was operated at 30 kV.

#### Data collection

Tilt-series data were collected using a Titan Krios G4 transmission electron microscope operating at 300 kV equipped with a Selectris X energy filter with a slit set to 10 eV and a Falcon 4i direct electron detector (Thermo Fisher Scientific) recording movies in Tiff format. A dose-symmetric tilt scheme using TEM Tomography 5 software (Thermo Fisher Scientific) was employed in the acquisition with a tilt span of ± 60°, covered by 2° steps starting at a 10º pre-tilt. Target focus was set for each tilt-series in a range of -2 μm to -4 μm in steps of 0.25 μm. The microscope was set to a magnification (64 000x) corresponding to a pixel size of 1.98 Å at the specimen level (corrected at the postprocessing step to 1.92 Å) and a nominal dose of ∼2.0 e^-^/Å^2^ per tilt image.

#### Tilt-series alignment and tomogram reconstruction

Data processing followed the workflow described in the TomoGuide^61^. See Supplementary Fig. 2 for processing work-flow. Using AreTomo3^62^, for 137 tilt series, raw tiff movies were automatically motion-corrected, their CTF estimated, tilt series aligned, and CTF-corrected tomograms reconstructed. Denoising was performed with Icecream or Isonet2 on tomogram pairs reconstructed from odd or even raw frames respectively^63,64^.

#### Subtomogram averaging

For particle picking, 3D template matching was carried out on bin4 CTF-corrected tomograms (7.64 Å/pixel) with 6° angular sampling in pyTOM-match-pick^65^ (Supplementary Fig. 2). Initial template matching (TM) was carried using a complex I reference obtained from sub-tomogram averaging in *C. reinhard-tii*^8^ and low-pass filtered to bin4 Nyquist (15.28 Å). This was initially performed on 9 tomograms. After particle extraction, classification and alignment in the tomography branch of RELION5^66^ using 2D stack particles, this yielded the initial 3D average of the I_1_III_2_ supercomplex from Phaeodactylum, that was used for another round of template matching, yielding more accurate picks. This was repeated another time, until our average reached bin4 Nyquist. A similar strategy was used for mitoribosomes, PDH and HSP60 (Supplementary Fig. 2).

Since respirasomes systematically went to highest resolution among all the targets, we used the output of RELION Bayesian Polishing job to generated realigned “polished” tomograms from the alignment of respirasome. A final round of TM on the polished tomograms job yielded 27639 I_1_III_2_ particles. Initial 3D classification without alignment revealed only a small fraction of false positive, and STA was performed with the remaining 27311 particles (Supplementary Fig. 5). Particles were aligned in RELION5 by iterative rounds of 3D refinement (using the --sigma_ang flag to strictly restrict RELION from deviating from the angular priors obtained from template matching) at bin4, 2 and 1, each followed by cycles of CTF Refinement and Bayesian Polishing, yielding the final map for I_1_III_2_ at 5.78 Å, followed by focused refinement on CI at 4.48 Å and CIII at 6.35 Å. From this set of aligned particles, 3D classification without alignment was performed at lower binning, initially with a large spherical mask, allowing to identify the main populations of I_1_III_2_ containing supercomplexes: dimer respirasome, crista respirasome, Oxphosome and IBM respirasome (Supplementary Fig. 5). These populations were then more accurately identified using specific masks covering supercom\plexes-specific regions. Iterative classification allowed to identify the boundary respirasome class, and particle lists comparison allowed to identify dimer respirasome supercomplexes associated with ATP synthase, a fusion of a dimer and oxpho-some, a class often visible by particle map back (Supplementary Fig. 7). Additionally, a class that could not be identified by 3D classification but by particle map back is “back-to-back” respirasomes (Supplementary Fig. 7a and c).

Alignment on CII and CIV was particularly challenging considering their size, relative positional heterogeneity, and predomi-nantly membrane-embedded regions. Starting from the aligned I_1_III_2_ particle set, focused 3D classification and refinement were then carried on the CII and CIV complexes and further refinement with the corresponding masks yielded maps at 10.48 Å and 9.87 Å, respectively (Supplementary Fig. 8). For CII two clear classes (Left and Right) were resolved, refined independently, then merged and refined together. For CIV, we initially attempted to 3D classify and refine all three CIVs independently and then recenter and merge the refined particles, but because CIVs are mostly membrane embedded and highly flexible relative to I_1_III_2,_ 3D refinement on all positions but CIV_”_(the most stable) failed. Hence, we only performed 3D refinement on CIV_”_. In parallel, 3D classification was performed on all three CIV_”_positions to assess their occupancy. This was performed in a iteration 3D classification with alignment with a mask covering each position. To determine the occupancy percentage shown in Fig. 2a, particle lists from CII and CIV classification were intersected with the supercomplex lists.

For the ATP synthase, we used the tetramer associated with the oxphosome as initial reference for ATP synthase detection by TM. Like the I_1_III_2_ detection, TM was repeated several times until the average reached bin4 Nyquist, and a single dimer (in-stead of the original dimer of dimer) was used for final detection, allowing to detect the different populations of ATP synthase (part or not of the oxphosome). Like the respirasomes, particles were aligned in RELION5 by iterative rounds of 3D refinement at bin4, and then bin2 until the dimer reached 12.13 Å (Supplementary Fig. 6). The dimers were refined with C2 symmetry, focused on the lateral stalks yielding 9.06 Å. In parallel, particles were symmetry expanded, and classification was performed to resolve the different catalytic states. 3D classification also revealed the ATP populations part of oxphosomes and the ones at the cristae tips. We thus obtained oxphosome classes both starting from ATP synthase of I_1_III_2_ template matching, with the ATP particles allowing to better resolve the ATP synthase part and I_1_III_2_ allowing the better resolve the respirasome part.

A similar general approach was used to resolve mitoribosomes, HSP60 and PDH (Supplementary Fig. 4).

### Membrane topography analysis

Membrane topography (shown in Fig. 2c) was measured on the subtomogram averages with custom python scripts. All maps were sampled at 5.76 Å/voxel in a common box, with the membrane normal along the box y axis. Membrane candidates were found without a density threshold by a Hessian-based sheet filter (σ = 15 Å). RANSAC planes fitted to these candidates were grown into quadric guide surfaces, and the surface covering the largest membrane area was kept. Along each column parallel to the normal, the bilayer mid-plane was first estimated from the two leaflet density peaks at a contour level of 0.0025. It was then refined without a threshold by cross correlating each column with the map’s own average bilayer profile (±50 Å, six it-erations). A column was accepted when the correlation was “ 0.4 and the two halves of the profile, one per leaflet, agreed on the mid-plane position within 10 Å (25 Å at the periphery, where the density fades); this excludes positions where protein density is fused to the membrane. The mid-surface was smoothed by Gaussian-weighted local plane fits (σ = 20 Å). Positions without detection, mainly beneath the complexes, were interpolated from the surrounding membrane. Membrane heights relative to the box center were reported within a circle of 23 nm radius around the box center. Curvatures κ_1_ and κ_2_ along two perpendicular axes were obtained by fitting h(s) = as^2^+bs+c to the detected membrane within 2 nm of each axis, with κ = 2a/(1+b^2^)^3/2^ at the circle center. Scripts are available at https://github.com/CryoFlo/cryoet-SpatialTools.

### Total membrane isolation and super-complexes separation

See Supplementary Fig. 10 for a schematic summary of the isolation. Phaeodactylum cells were pelleted down from 1-2 L of cell culture grown in ESAW medium by centrifugation at 2000 g for 10 min at room temperature. Pellet was resuspended in 50 mL of Isolation Buffer (IB: 0.3 M mannitol, HEPES-KOH 30 mM, EDTA 2 mM, MgCl_2_ 5 mM, KCl 10 mM, MnCl_2_ 1 mM, PVP 1 w/v, pH adjusted to 7.5). Resuspended cells were disrupted in a high-pressure homogenizer (EmulsiFlex-C5, Avestin, Canada) at 900-1200 bar. Monitoring of fractions under light microscope showed that three cycles were enough for effective cell lysis. After lysis, the cells were diluted with cold IB to 240 mL and homogenized with glass Dounce potter using loose piston to disperse possible residual membrane aggregates. The resulting solution was centrifuged at 500 g for 6 min at 4°C twice to pellet cell debris. Resulting supernatant with organellar extract was pelleted at 15k g for 15 min at 4°C, and the pellet was resuspended in 50 mL of cold washing buffer (WB: mannitol 0.3 M, HEPES-KOH (pH 8) 30 mM, EDTA (pH 8) 2 mM, MgCl_2_ 5 mM, KCl 10 mM, MnCl_2_ 1 mM, pH adjusted to 7.5 by HCl/KOH) and centrifuged at 14k g for 15 min, which was repeated twice for washing out residual PVP from IB buffer. The crude membrane extract was then pelleted, frozen into liquid N_2_ and stored at -80°C.

For separation of mitochondrial membrane complexes, crude membrane fractions were resuspended in 2 mL mitochondria lysis buffer (LB: 20mM HEPES-KOH, pH 7.5, 75 mM KCl, 1 mM DTT, 5% Digitonin), mixed by pipetting and incubated for 30 min on a rotating wheel at 4°C. The lysate was clarified by centrifugation at 14k g for 10 min at 4°C and the supernatant was homogenized in glass Dounce potter using tight piston. 1 mL of the lysed membrane fraction was then loaded on a 10-40% continuous sucrose gradient in the same buffer supplemented with 0.1% digitonin and centrifuged for 15h at 31k g, 4°C, in a SW41 rotor. Fractions were automatically collected using a BioComp gradient fractionator, pooled, and pelleted by centrifugation for 3 h at 500k g, 4°C, in a TLA-120.2 rotor.

### Sample digestion and proteomics analysis

Fractions of Oxphos complexes were reconstituted in lysis buffer (5% SDS, 10mM TCEP, 0.1 M TEAB), incubated for 10 min at 95°C, cleared by centrifugation and digested using S-Trap™ micro spin columns (Protifi) according to the manufacturer’s instructions. Shortly, 12 % phosphoric acid was added to each sample (final concentration of phosphoric acid 1.2%) followed by the addition of S-trap buffer (90% methanol, 100 mM TEAB pH 7.1) at a ratio of 6:1. Samples were mixed by vortexing and loaded onto S-trap columns by centrifugation at 4000 g for 1 min followed by three washes with S-trap buffer. Digestion buffer (50 mM TEAB pH 8.0) containing sequencing-grade modified trypsin (1/25, w/w; Promega, Madison, Wisconsin) was added to the S-trap column and incubate for 1h at 47 °C. Peptides were eluted by the consecutive addition and collection by centrifugation at 4000 g for 1 min of 40 ul digestion buffer, 40 ul of 0.2% formic acid and finally 35 ul 50% acetonitrile, 0.2% formic acid. Samples were dried under vacuum and stored at -20 °C until further use. Dried peptides were resuspended in 0.1% aqueous formic acid and subjected to LC– MS/MS analysis using a Orbitrap Fusion Lumos Mass Spectrometer fitted with an EASY-nLC 1200 (both Thermo Fisher Scientific) and a custom-made column heater set to 60°C. Peptides were resolved using a RP-HPLC column (75μm × 36cm) packed in-house with C18 resin (ReproSil Saphir 100 C18, 1.5 um resin; Dr. Maisch GmbH) at a flow rate of 0.2 ul/min. The following gradient was used for peptide separation: from 2% B to 8% B over 5 min to 25% B over 40 min to 35% B over 15 min to 95% B over 2 min followed by 10 min at 95% B. Buffer A was 0.1% formic acid in water and buffer B was 80% acetonitrile, 0.1% formic acid in water.

The mass spectrometer was operated in DIA mode. MS1 scans were acquired in the Orbitrap in centroid mode at a resolution of 120,000 FWHM (at 200 m/z), a scan range from 390 to 910 m/z, normalized AGC target set to 250 % and a maximum ion injection time of 50 ms. MS2 scans were acquired in the Orbitrap in centroid mode at a resolution of 15,000 FWHM (at 200 m/z), precursor mass range of 400 to 900, quadrupole isolation window of 16 m/z with 1 m/z window overlap, a scan range from 145 to 1450 m/z, normalized AGC target set to 2000 % and a maximum ion injection time of 22 ms. Peptides were fragmented by HCD (Higher-energy collisional dissociation) with collision energy set to 33 % and one microscan was acquired for each spectrum. The acquired raw files were searched using DIA-NN (v.2.2.0) in “library-free search” mode against a Phaeodactylum tricornutum database (consisting of 10484 protein sequences downloaded from Uniprot on 20250113), spiked with the amino acid sequences of F1DGQ2, F1DGQ3, F1DGM8, F1DGM9, F1DGP3, F1DGM2, F1DGM6, F1DGQ1, F1DGM3, F1DGQ5, F1DGM1, A0A8J9TDK6, F1DGQ4, F1DGN0, F1DGM7, F1DGM5, F1DGN2, F1DGN8, F1DGQ0 and 393 commonly observed contaminants using default library-free search settings with the following modifications: 1) oxidation of methionine and protein N-terminal acetylation were set as variable modifications, 2) Protein inference was set to “Protein names (from FASTA)”.

### Integrative Model building and refinement

See Supplementary Fig. 11 & 12 for initial AlphaFold3 and final models. To build the models of complex I to V, we used an integrative approach to identify the protein subunits of the different complexes. Conserved core subunits were identified by querying the proteomes of *Phaeodactylum tricornutum, Thalas-siosira pseudonana*, and related diatom species using BlastP^33^ and Foldseek^34^ for sequence and structural homology, respectively. Most of the core subunits were identified this way by sequence or structural homology. These proteins were then used as markers to track the different complexes by MS and identify potential species-specific subunits or subunits that we failed to identify by sequence and structural homology (Supplementary Fig. 10). The identified proteins were then predicted using Al-phaFold3^35^ and Protenix^67^ multimer, either on local installations and/or web-based platforms on either the whole complexes or only portions of them if too large (notably for complex I). The initial models were then refined, by trimming or completing the missing protein parts, by comparing the predicted models against the subtomogram averages and monitoring the pLDDT scores. The primary sequence database was derived from Uni-Prot, but in some case we had to go back to the reference genomes^22,68^ to curate the annotations. For the unknown component (e.g the FAS domain on complex I), were identified through an iterative screening process where known components of the matrix arm were predicted together with the MS-identified candidates, until a predicted complex would fit the density. Some protein subunits (all conserved mitochondria-specific) could not be identified and were either replaced by polyA peptide or protein from other species, see Supplementary Fig. 11 & 12.

Initial multimeric predictions were docked into their subtomogram averages via rigid-body fitting in UCSF ChimeraX^69^. Again, models were subsequently optimized by trimming or adding disordered termini. Once we obtained the final predicted models, they were then manually inspected and refined in COOT^70^. For complex I and III, visible conserved ligands, such as iron sulfur clusters and hemes were added. The different parts of the model were then automatically refined in PHENIX using the phenix.real_space_refine tool. For higher-order supercomplex architectures, they were assembled by rigid-body fitting the refined individual complex models into the corresponding supercomplexe map densities. The model geometry was validated using MolProbity^71^ and is summarized in Supplementary Table 1.

### Complex I FAS-like domain sequences

Candidate homologues of ND-PCKL, ND-ACSL and ND-TERL were identified in selected eukaryotic proteomes using sequence searches with DIAMOND v2.1.15 in ultra-sensitive mode and profile searches with HMMER v3.4. Queries included the *Phaeodactylum tricornutum* subunits, corresponding *Euglena gracilis* sequences from PDB 8J9J, the *Tetrahymena thermophila* ACSL subunit and active enzyme-family comparators. Representative sequences were selected using sequence and profile similarity, with reciprocal searches used to assess selected candidates. Each protein family was aligned independently using MAFFT v7.526. Final datasets comprised 18 PCKL, 19 ACSL and 21 TERL sequences, including active comparators P22259, P9WQ37 and Q8WZM3, respectively (listed in Supplementary Table 3). Alignments are displayed in taxonomic order, with exact identity to the corresponding active comparator indicated in black.

### Molecular dynamics

Each protein complex was converted to a Martini 3^48^ representation using martinize2^72^, merging all chains and using an elastic network of 700 kJ mol^-1^ nm^-2^ with default protonation states. Each complex was then embedded into a symmetric membrane of POPE:POPC:CDL2 (4:2:1), based on previous studies^10^, using insane^73^ and the systems were solvated. 150 mM NaCl was added after neutralising the system. Energy minimisation was performed using the steepest descents method. A timestep of 20 fs was used for production simulations and performed in an NpT ensemble at 310 K and 1 bar. Temperature coupling was performed with the Velocity rescaling algorithm^74^, with protein, lipids and solvent & ions coupled separately. The temperature coupling constant was set at 1 ps. The Parrinello-Rahman algorithm^75^ was used for pressure coupling with a τp = 12.0 ps in a semi-isotropic manner. Van der Waals interactions used a single cutoff of 1.1 nm, and the reaction-field algorithm was used for electrostatic interactions with a cutoff of 1.1 nm^76^. The settings verlet-buffer-tolerance = -1 and rlist = 1.35 nm were used to prevent possible neighbour-list artefacts^77^. All simulations were performed with GROMACS 2024.3^78^. For each complex, five independent simulations of 5 μs were conducted.

For single complex V dimer simulations, a reduced version was used with the soluble region truncated ∼4 nm from the bilayer. Tetramer simulations had lipid number asymmetry of ∼10 % to allow membrane bending, with one simulation of 1 μs performed due to the system size^56,79^.

Lipid density analysis was performed using a previously published script^80^ and occupancy was measured using PyLipID^81^. Membrane thickness and membrane curvature were measured using an in-house script utilising MDAnalysis^82,83^ and matplotlib^84^.

### Figure preparation

All figures were prepared using UCSF ChimeraX, developed by the Resource for Biocomputing, Visualization, and Informatics at the University of California, San Francisco, with support from National Institutes of Health R01-GM129325 and the Office of Cyber Infrastructure and Computational Biology, National Institute of Allergy and Infectious Diseases^85,86^. For tomogram views with map-backs particle map back (Fig.1 and 2 and Supplementary Fig. 7), the ArtiaX plugging was used^87^ and membrane segmentations for visualization were automatically performed using MemBrain-seg and manually refined^88^.

## Supplementary materials

The manuscript also includes Supplementary Figures 1 to 15, Supplementary Tables 1 to 3 and Supplementary Videos 1 and 2.

## Notes

### Competing Interest Statement

The authors have declared no competing interest.

