## Supplementary figures 1 to 15 for "Native architecture of a mitochondrial oxphosome"

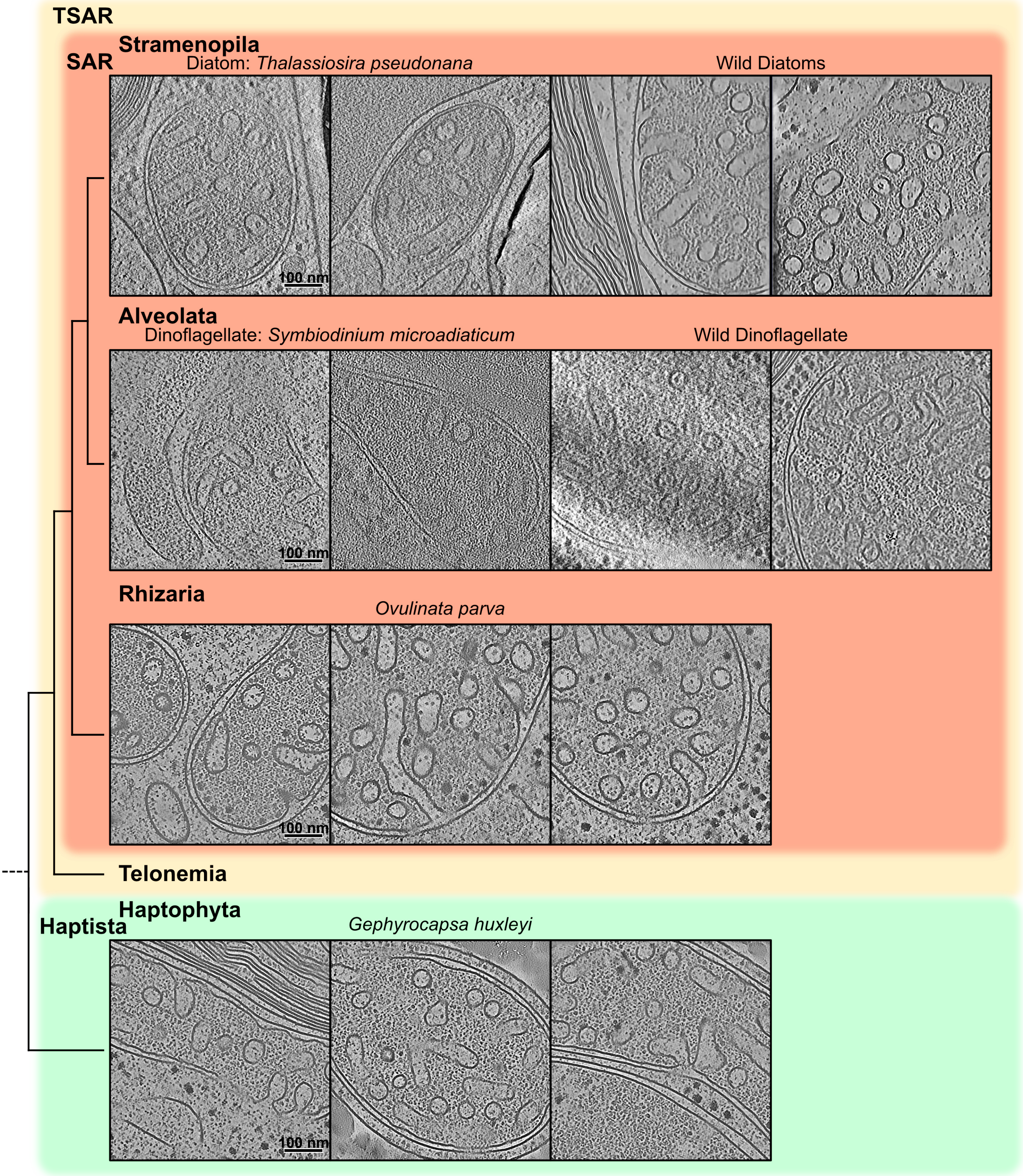


**Supplementary Figure 1 :** Tomogram gallery of tubular cristae mitochondria

Representative tomograms highlighting mitochondrial structures across key lineages within the TSAR and Haptista supergroups. The SAR clade is represented by Stramenopila, featuring the cultured diatom *Thalassiosira pseudonana* and wild a diatom (together with Schizochytrium presented in Fig. 1 and *Phaeodactylum tricornutum* whose mitochondrial molecular architecture is described in detail throughout the manuscript); Alveolata, showing the dinoflagellate *Symbiodinium microadriaticum* and a wild dinoflagellate; and Rhizaria, illustrated by *Ovulinata parva*. The Haptista supergroup is represented by the Haptophyta species *Gephyrocapsa huxleyi* (formerly Emiliana). While Telonemia is included in the phylogenetic framework, no tomograms are displayed for this group; however, Telonemia was also shown to have tubular cristae by classical EM in Tikhonenkov et al[18]. The wild diatoms and dinoflagellate were sampled in Kristineberg in Sweden. All the tomograms are shown at the same scale.

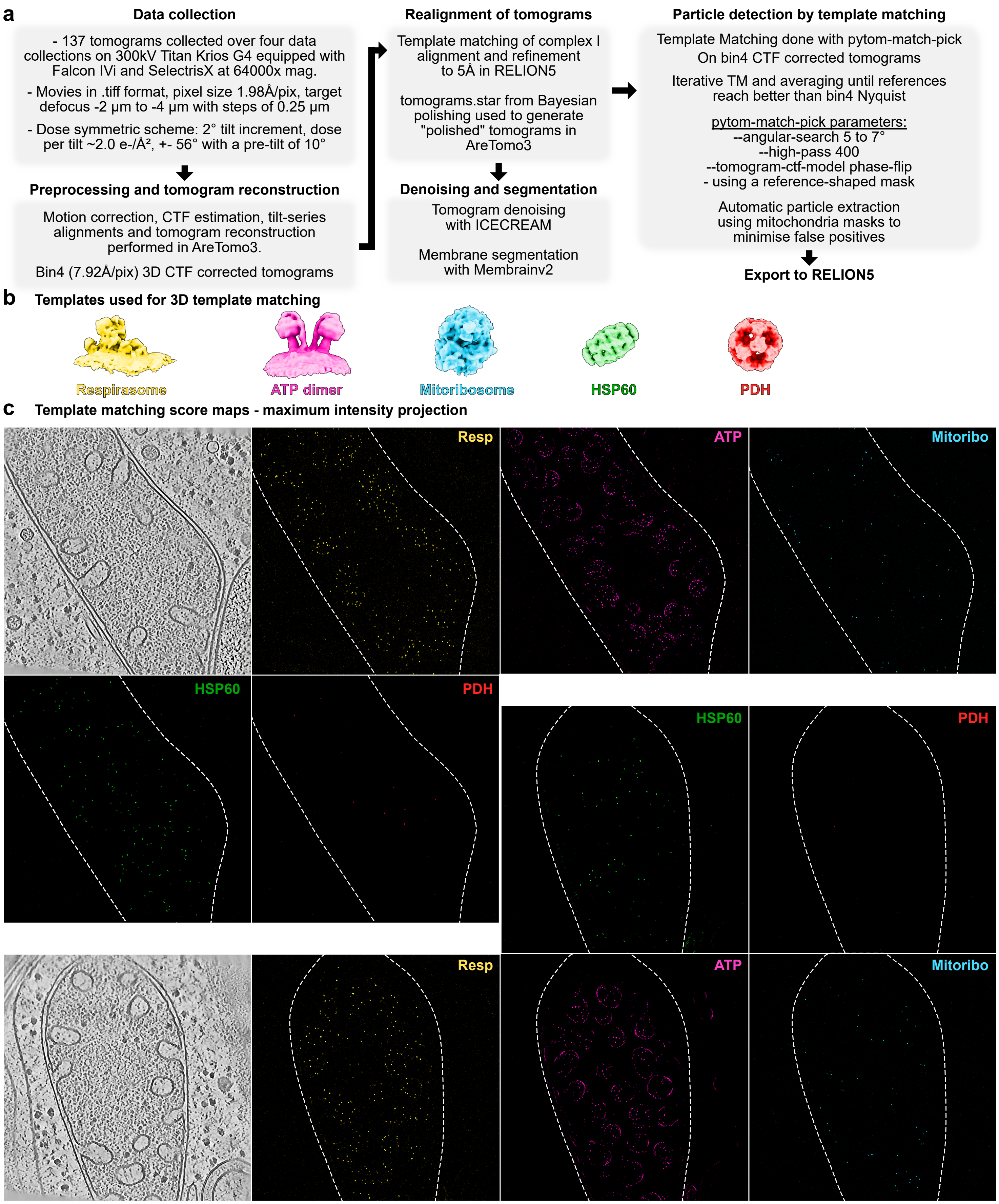


**Supplementary Figure 2 :** Tomography data processing summary

**a**, Workflow for cryo-ET data acquisition, preprocessing, tomogram reconstruction, denoising, membrane segmentation, and particle detection. A total of 137 tomograms were collected on a 300 kV Titan Krios G4 microscope equipped with a Falcon 4i detector and Selectris X energy filter. Preprocessing and 3D CTF-corrected tomogram reconstruction were done in AreTomo3 at bin4 (7.92 Å/pixel). Particle detection was performed using 3D template matching in pytom-match-pick on bin4 CTF-corrected tomograms with iterative averaging, followed by automatic particle extraction using mitochondrial masks and export to RELION5. **b**, Templates used for 3D template matching: respirasome, ATP synthase dimer, mitoribosome, HSP60, and pyruvate dehydrogenase (PDH) complex. **c** Maximum intensity projections of template matching score maps shown for two representative tomograms across the five target complexes. Each colored dot represents a high cross correlation score, and thus the position of the target complex.

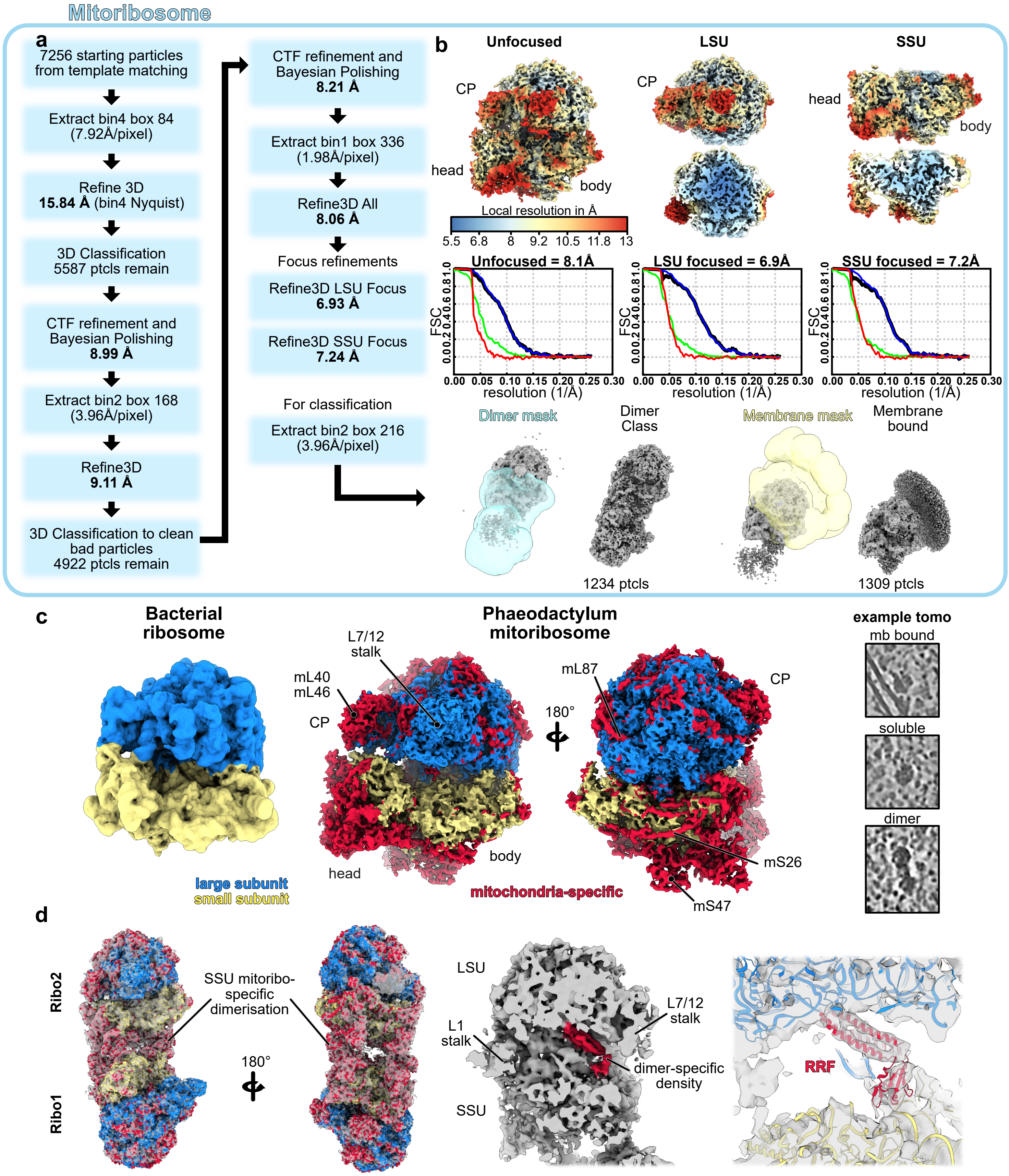


**Supplementary Figure 3 :** STA Processing and details of the Phaeodactylum mitoribosome

**a**, STA image-processing pipeline used to obtain the maps of the Phaeodactylum mitoribosome monomer, dimer, and membrane-bound classes. **b**, Gold-standard FSC curves are plotted with resolution reported at the FSC = 0.143 criterion. Local-resolution estimates are displayed on the corresponding maps using a common color scale (in Å). **c**, Comparison of the Phaeodactylum mitoribosome with a bacterial ribosome structure (EMD-43929). Large subunits are colored in blue, small subunit in yellow and additional protein and rRNA domains are highlighted in red. Identifiable mitoribosome-specific proteins are annotated. CP for Central Protuberance. **d**, Structural insights into hibernating mitoribosome dimers. Dimer interface mediated by mitochondria-specific domains of the SSU. Cutaway view of the subunit interface reveals an additional density unique to the dimer class. Atomic model fit (PDB: 6ERI) revealing that this dimer-specific density likely corresponds to Ribosome Recycling Factor (RRF), characteristic of a stalled or hibernating ribosomal state.

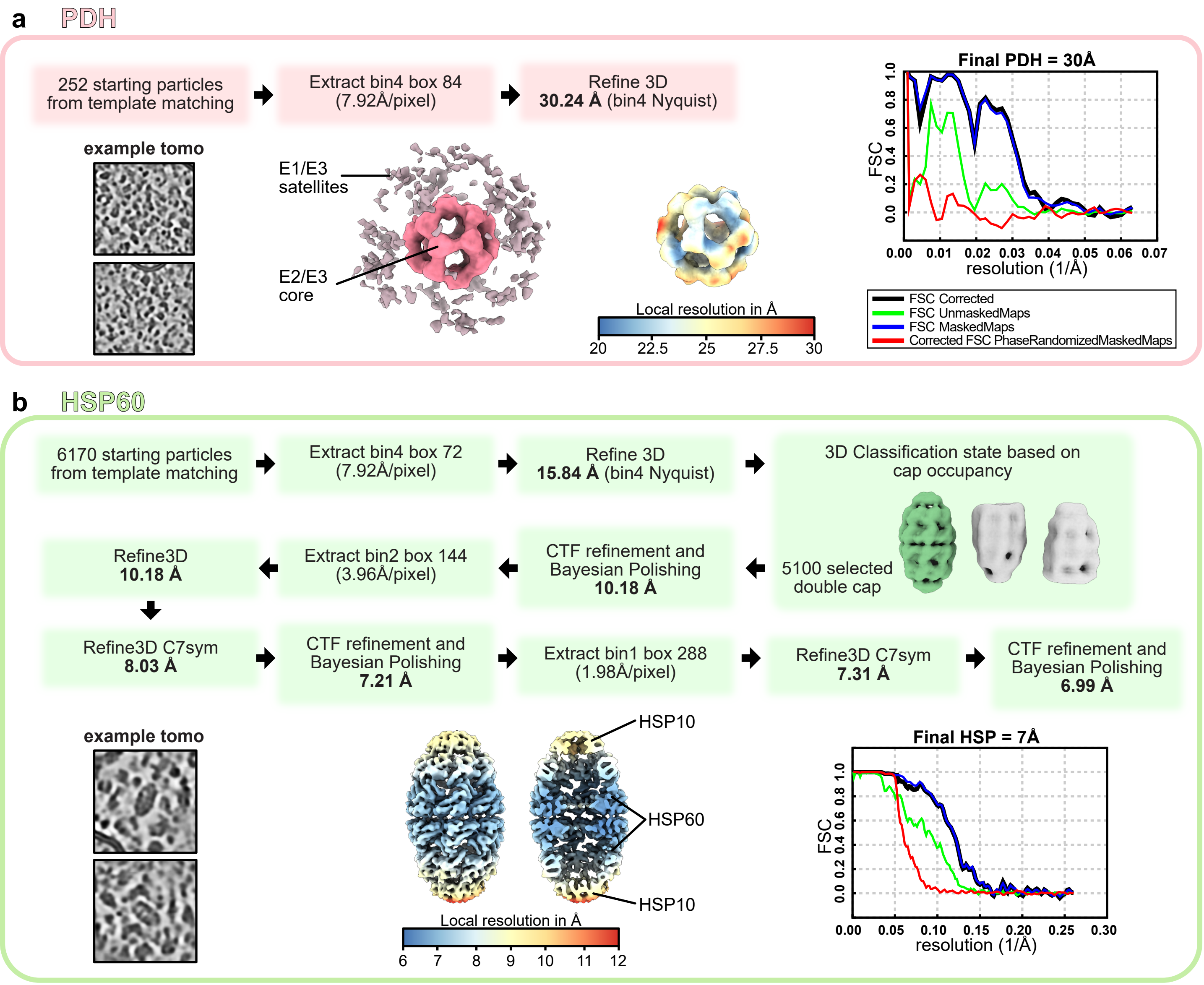


**Supplementary Figure 4:** STA processing summary for PDH and HSP60

**a**, and **b**, Workflows used to reconstruct the PDH (**a**) and HSP60 (**b**) complexes. Key structural components for each complex are annotated on their final 3D density. Local resolution estimates (in Å) are mapped onto the corresponding density maps alongside colored scale bars. Gold-standard Fourier shell correlation (FSC) curves are plotted for both reconstructions, with final overall resolutions reported using the FSC = 0.143 criterion.

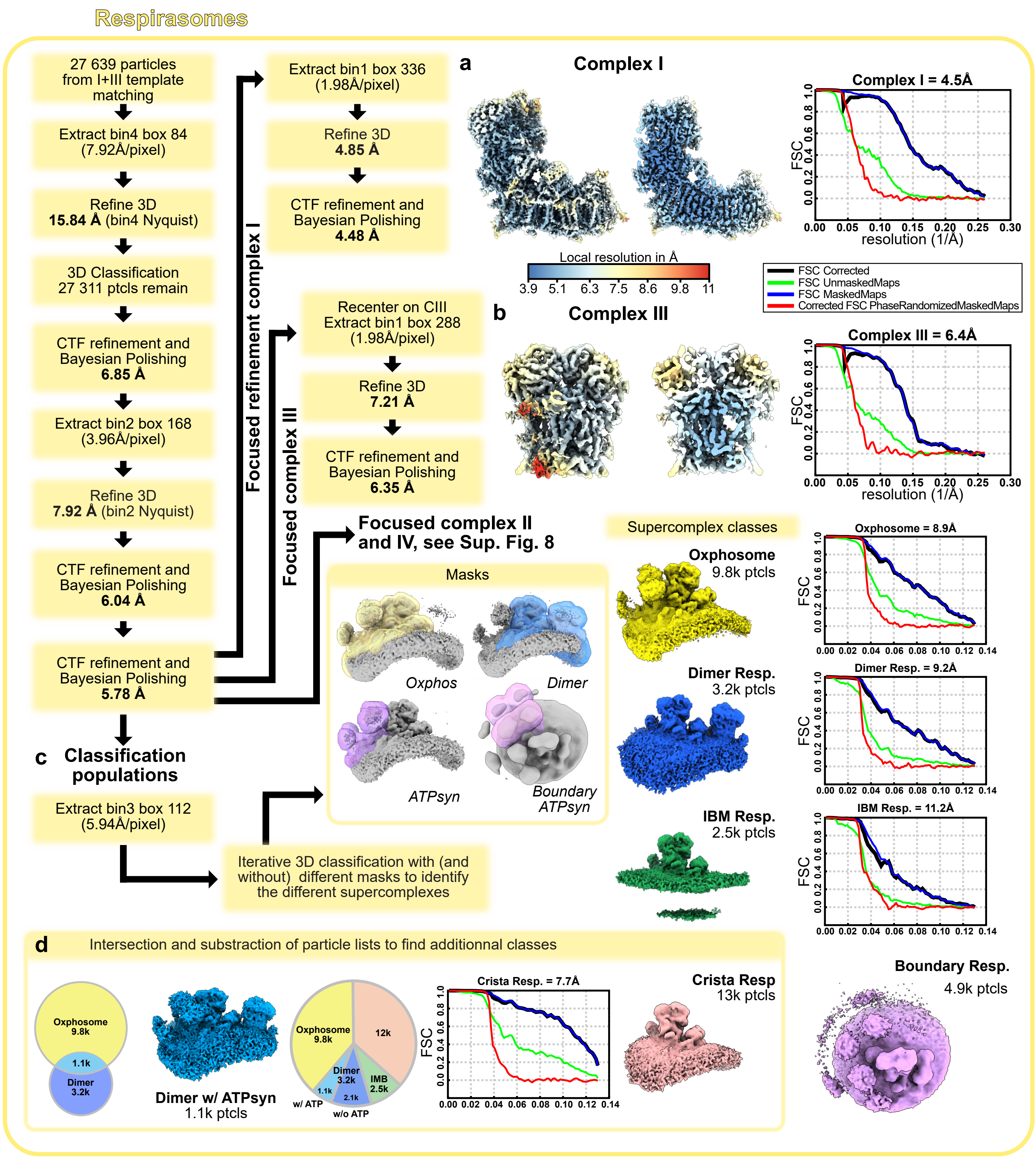


**Supplementary Figure 5 :** STA processing summary for respiratory supercomplexes

Following an initial common processing workflow focused on $I_1III_2$ core, the workflow diverges several branches. **a**, Focused on CI. **b**, Focused on CIII. For CII and CIV, see Supplementary Fig. 8. **c**, Iterative 3D classification to find different supercomplex populations: Oxphosome, Dimer respirasome and IBM respirasome and the boundary respirasome. **d**, Additional classes were found by particle list intersection and subtraction. For the final high-resolution reconstructions in **a** and **b**, local resolution estimates (in Å) are mapped onto the corresponding density maps alongside colored scale bars, and gold-standard Fourier shell correlation (FSC) curves are plotted.

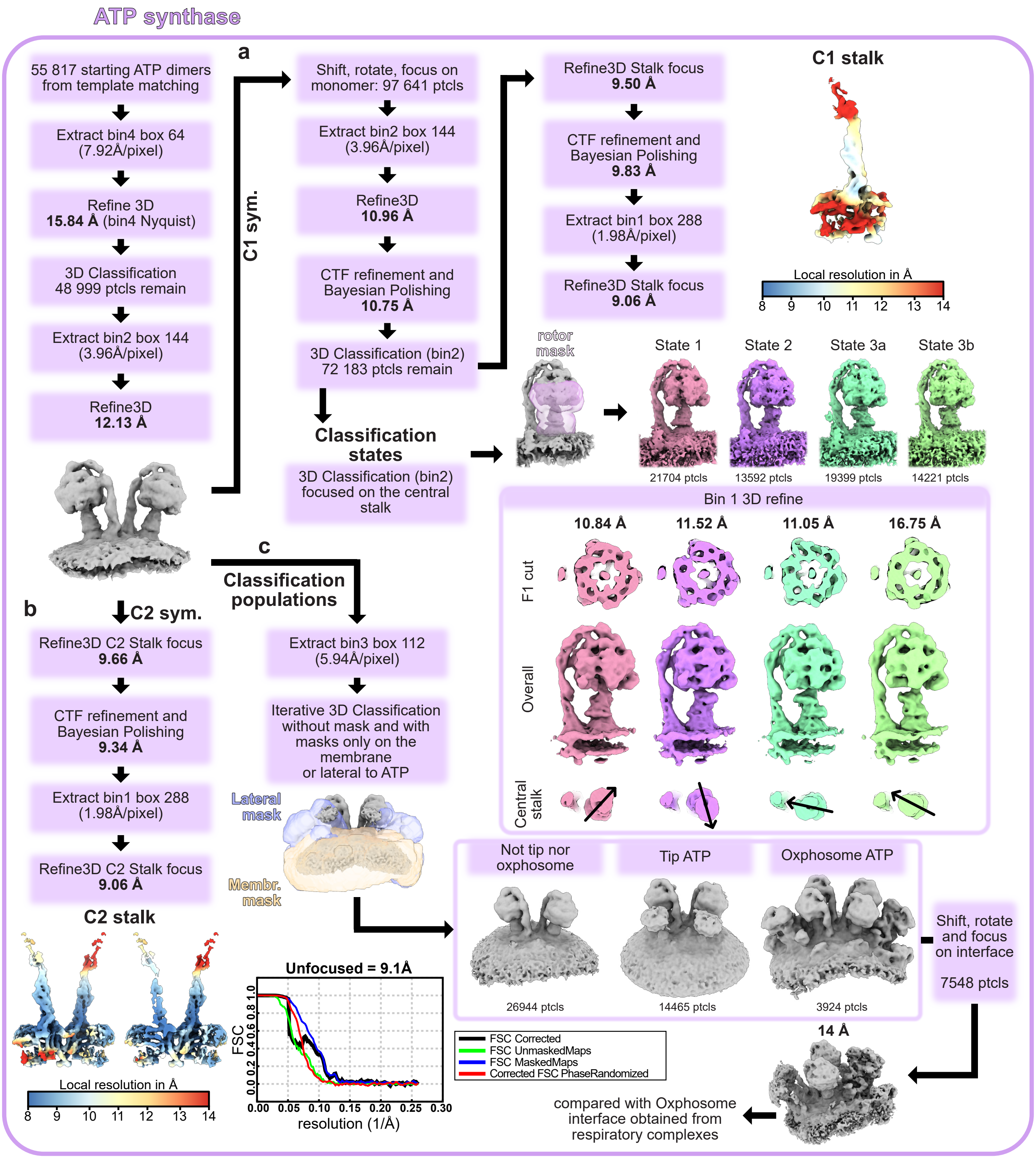


**Supplementary Figure 6:** STA processing summary for ATP synthase

Following an initial common processing workflow for ATP synthase dimers, the pipeline diverges into three branches. **a**, Focused monomer refinement with C1 symmetry. 3D classification focused on the central stalk resolved four distinct functional rotary substates (States 1, 2, 3a, and 3b). Rotation of the central stalk is indicated with an arrow. **b**, Dimer refinement applying C2 symmetry and focused on the lateral stalk, which yielded improved structural resolution compared to the C1 consensus refinement. **c** Iterative 3D classification of dimer subpopulations. Utilizing specific membrane and lateral masks, distinct higher-order assembly classes are resolved, including «Tip ATP» and «Oxphosome ATP» populations. From the Oxphosome class, alignment was performed on the CI-ATP interface, as done with the Oxphosome obtained from $I_1III_2$ particles, but the interface from $I_1III_2$ complexes yielded a better reconstruction. For the final high-resolution reconstructions in **a** and **b**, local resolution estimates (in Å) are mapped onto the corresponding density maps alongside colored scale bars, and gold-standard Fourier shell correlation (FSC) curves are plotted.

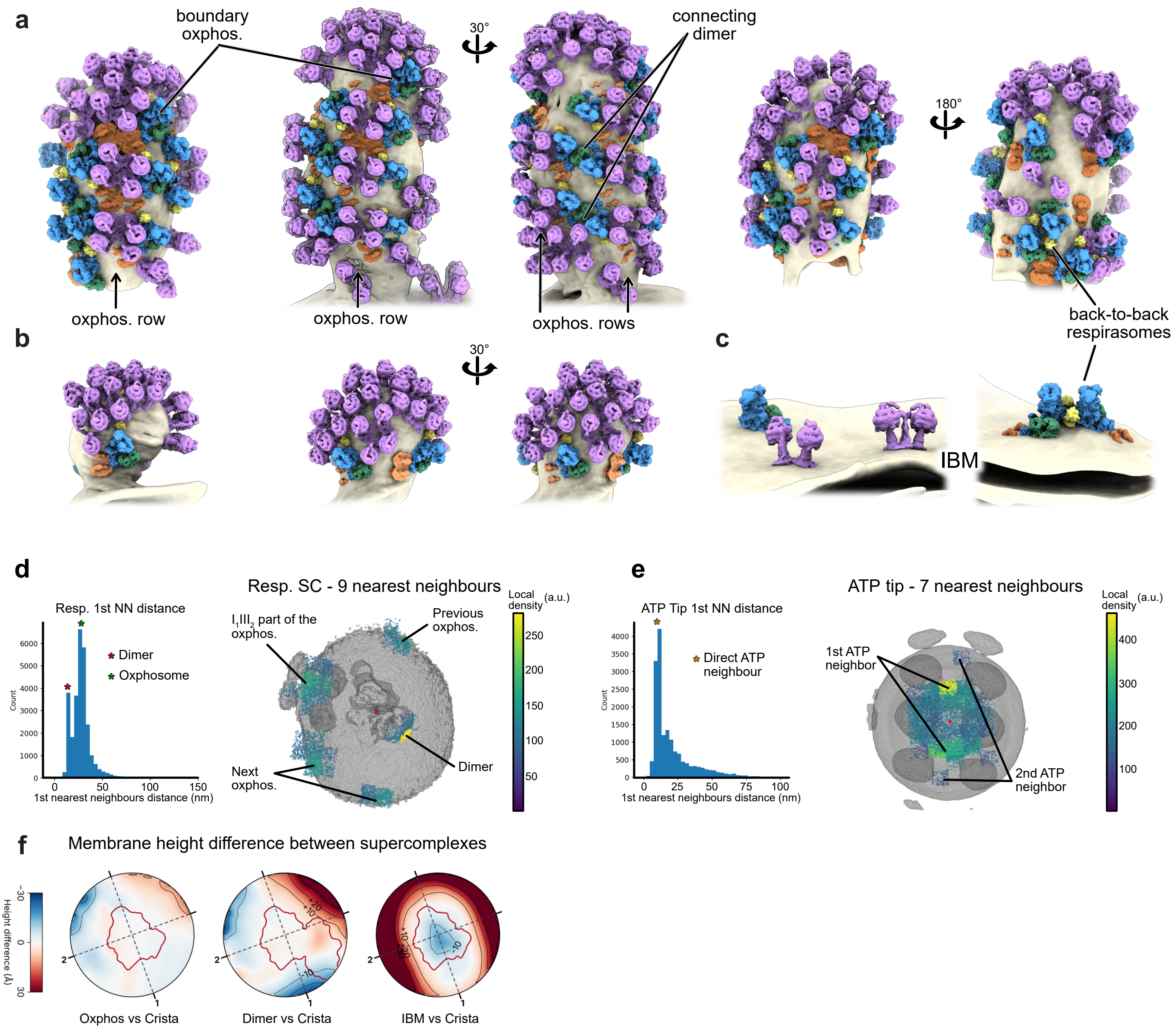


**Supplementary Figure 7:** Higher order complexes organization

**a-c**, Similarly to Fig. 2a, additional cristae are shown, highlighting the occurence of oxphosome rows, the presence of respirasome dimers connecting oxphosome rows, **b**, initial binding of boundary oxphosomes on nascent cristae, as well as **c**, isolated complexes on the IBM, and both present on cristae and IBM "back-to-back" respirasomes that we were not able to obtain by 3D classification. **d** and **e**, Nearest-neighbour distance distributions for all respiratory SC (b) and ATP synthase tip (c) particles. **d**, Left: distribution of nearest-neighbour distances; stars mark the respirasome dimers (red) and the adjacent $I_1III_2$ part of the same oxphosome (green). Right: positions of the 9 nearest neighbours of each particle, shown in the frame of the central particle (red dot) over its subtomogram average shown from the matrix view (grey) and coloured by local density (a.u.). Points with value < 90 a.u. are hidden for clarity. The clusters reflect the regular higher-order organization shown in **a**, and their identity is indicated. **e**, As in **d**, with the 7 nearest neighbours of ATP synthase tip particles; the star marks the directly adjacent ATP synthase. Points with value < 110 a.u. are hidden for clarity. Here, past the directly adjacent ATP synthase, no clear order emerges. **f**, Membrane height differences between the main supercomplex classes shown in Fig. 2c. Height of the membrane mid-surface in each map minus that of the crista respirasome, shown in the same orientation as the height maps. Red and blue mark regions where the membrane is higher or lower than in the crista respirasome (±30 Å; contours every 10 Å, negative values dashed). The red outline marks the footprint of the fitted model, and dashed lines 1 and 2 are the axes used for $\kappa_1$ and $\kappa_2$. The Oxphosome and crista respirasome have nearly identical membrane shapes, whereas the flat IBM membrane departs strongly from the curved crista membrane. In the dimer respirasome, the membrane is tilted relative to the crista respirasome: lower next to the second copy and higher on the opposite side.

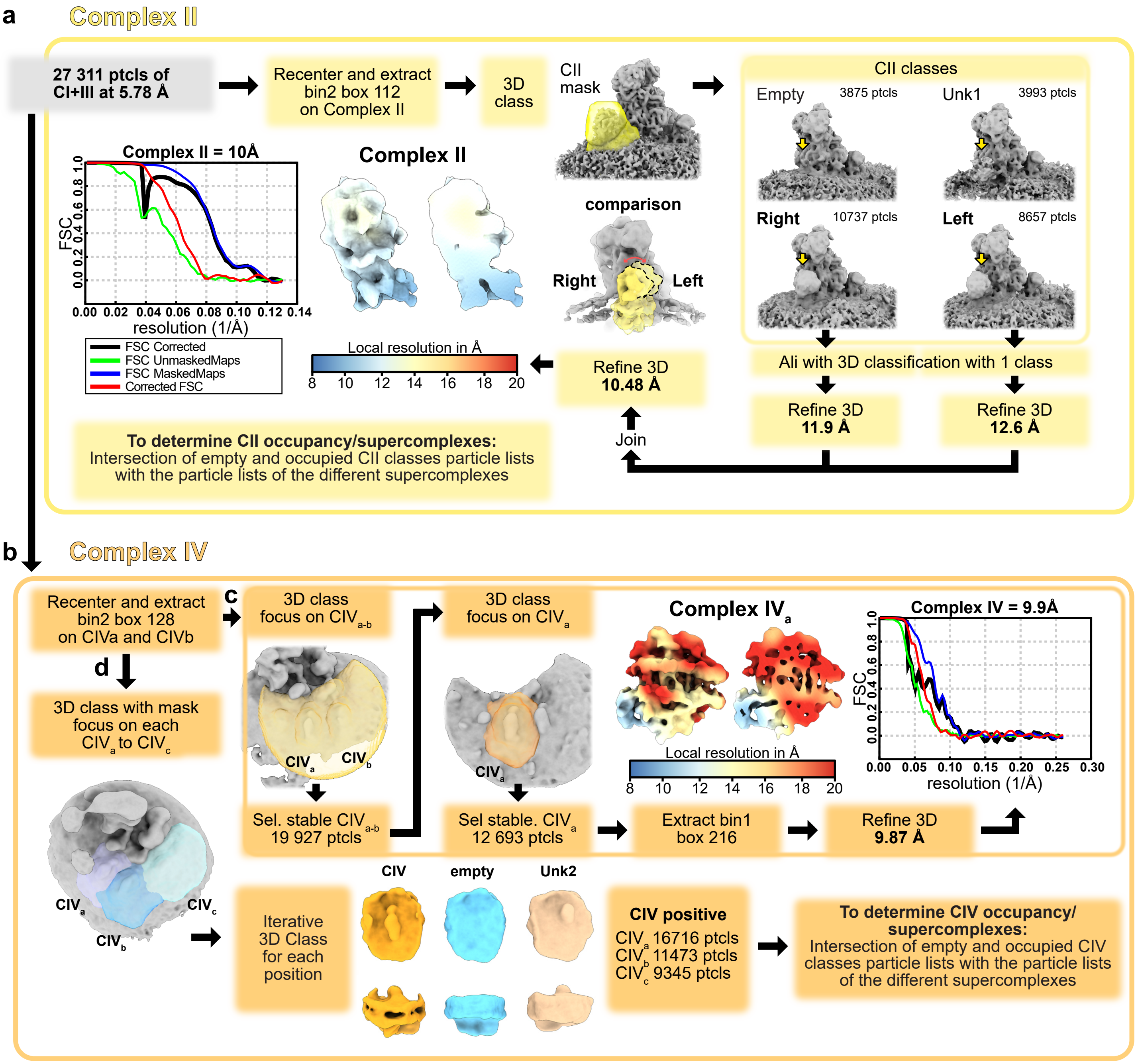


**Supplementary Figure 8:** Refinement and flexible association of complex II and IV

Starting from the aligned $I_1III_2$ particle set from Supplementary Fig. 5, the processing workflow branches into dedicated pipelines for CII (**a**) and CIV (**b–d**). **a**, Subtomogram averaging and classification pipeline for complex II. Particles were recentered, extracted, and subjected to 3D classification using a CII mask to isolate distinct structural classes. Two clear classes (Left and Right) were resolved, plus an unknown (Unk) and empty classes. Comparison of the Left and Right classes is shown. Both were individually refined, then re-combined underwent 3D refinement to yield a final reconstruction. **b–d**, Processing and focused classification strategies for complex IV ($CIV_a$ and $CIV_{a-c}$). **c**, To obtain the best CIV map, iterative focused 3D classifications were performed first with a mask covering $CIV_{a-b}$, then with a mask only on $CIV_a$. Particles were re-extracted at bin1 and refined to a final high-resolution map. **d**, For CIV occupancy, we used a mask on each CIV position. Iterative 3D classification was performed for each position on the basis of CIV occupancy. Typical CIV positive or empty classes are shown from top and side views (positional heterogeneity for $CIV_a$ and $CIV_b$ are shown in Supplementary movie 2), as well as Unk2 which is a recuring class only found at the $CIV_b$ position, that might correspond to a CII occupancy class. For both CII and CIV (**a** and **b**), their respective occupancy per supercomplex, shown in Fig. 2b, was determined by cross the classification particle lists with the supercomplex particle lists. For the final reconstructions, local resolution estimates (in Å) are mapped onto the corresponding density maps alongside colored scale bars, and gold-standard Fourier shell correlation (FSC) curves are plotted.

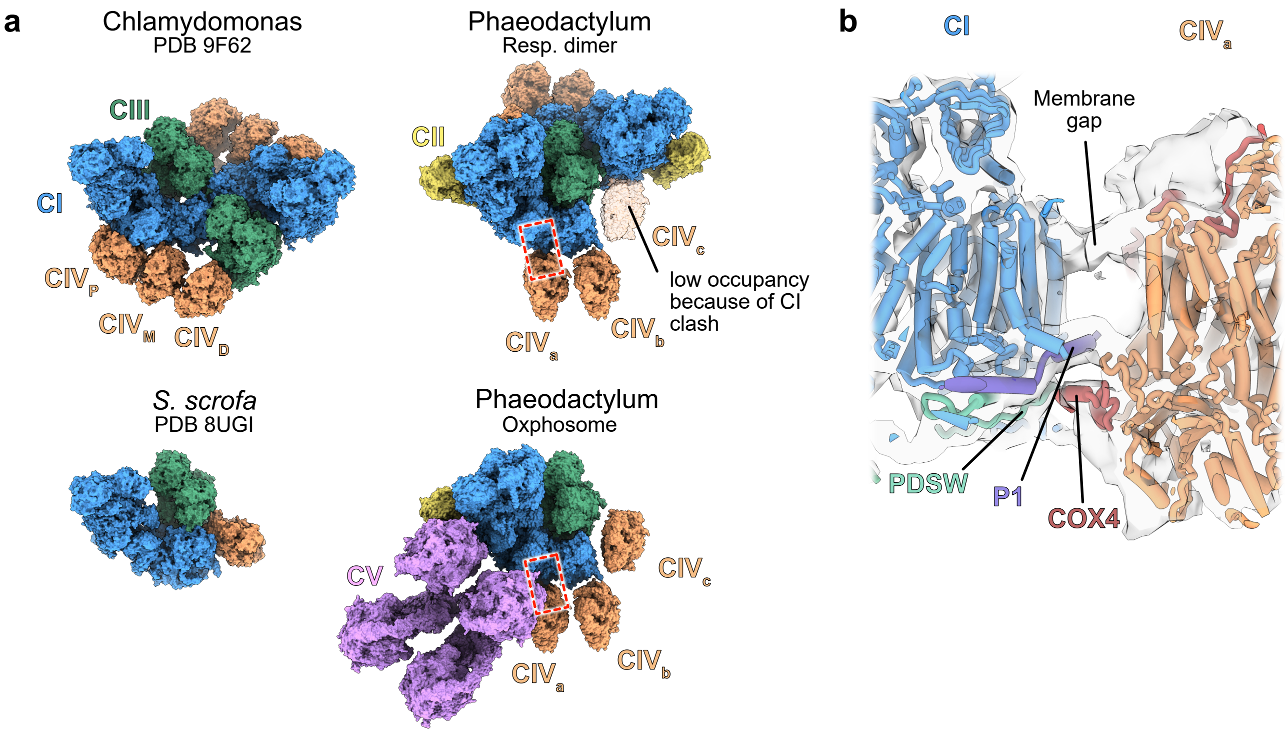


**Supplementary Figure 9:** Structural comparison and interactions of complex IV across mitochondrial supercomplexes

**a**, Structural comparison of CIV locations within mitochondrial supercomplexes from *C. reinhardtii* (PDB 9F62), *S. scrofa* (8UGI), and Phaeodactylum respirasome dimer and oxphosome with complex I (CI) in blue, complex II (CII) in yellow, complex III (CIII) in green, complex IV (CIV) in orange, and complex V (CV) in pink. In Phaeodactylum, the position of $CIV_a$ is analogous to $CIV_M$ in Chlamydomonas, whereas $CIV_c$ corresponds to the canonical mammalian CIV position. The position typically occupied by $CIV_P$ in Chlamydomonas is occupied by CV in the Phaeodactylum oxphosome. Within the respirasome dimer, $CIV_c$ exhibits substantially lower occupancy than other Phaeodactylum supercomplexes, likely due to a steric clash with the second CI. **b**, Zoomed-in view of the CI - $CIV_a$ interface (indicated by the red dashed rectangle in **a**), representing the only CIV copy that forms direct protein-protein interactions with CI. Key interacting subunits are annotated.

**a**

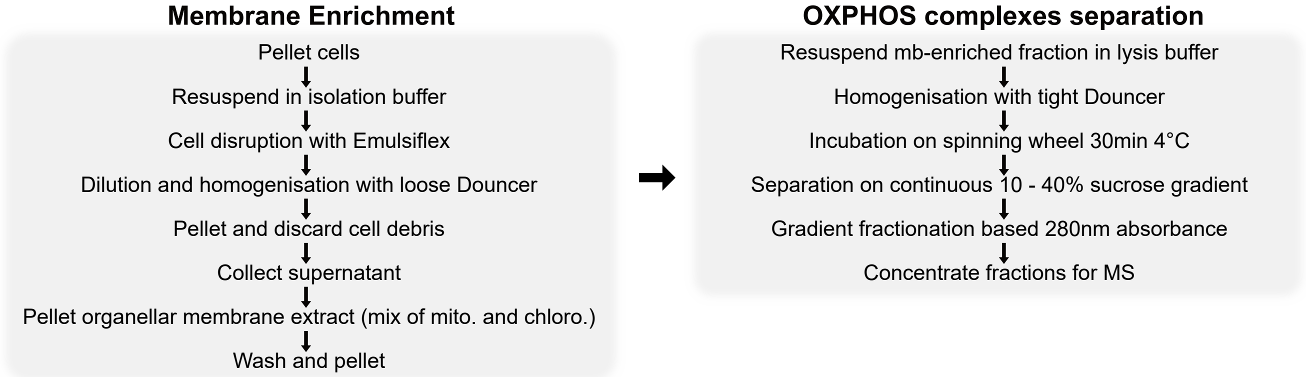


**b**

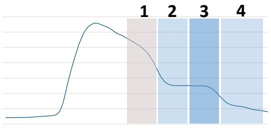


CI

| Matrix | Phaeo ID | Length | Abundance |
|---|---|---|---|
| 51kDa | B5Y4V1 | 493 | |
| 75kDa-Nter | F1DGQ2 | 205 | |
| Nad7m/ND7 | F1DGM8 | 398 | |
| PSST | B7GES5 | 166 | |
| TYKY | B7G4H7 | 163 | |
| 24kDa | B7G964 | 260 | |
| B17.2 | B7FVX2 | 219 | |
| B14 | B7S480 | 123 | |
| B8 | B7FUM1 | 262 | |
| B13 | B7GA70 | 194 | |
| 18kDa | B7G8A8 | 314 | |
| 13kDa | B7G589 | 127 | |
| 39kDa | B7FQ48 | 397 | |
| B14.5a * | B7G8T7 | 278 | |
| SDAP1 | B7FRC2 | 127 | |
| SDAP3 | B7FRC2 | 127 | |
| ND-PCKL * | B7FYE8 | 575 | |
| ND-ACSL * | B7G0K8 | 285 | |
| ND-TERL * | B7G395 | 500 | |
| **Membrane proximal** | **Phaeo ID** | **Length** | **Abundance** |
| Nad1m/ND1 | F1DGP3 | 328 | |
| Nad2m/ND2 | F1DGM2 | 546 | |
| Nad4Lm/ND4L | F1DGQ1 | 106 | |
| Nad6m/ND6 | F1DGM1 | 200 | |
| B14.7 | B7G570 | 160 | |
| NUXM | B5Y404 | 159 | |
| B16.6 | B7G3W2 | 136 | |
| CA-Like | B7GE24 | 201 | |
| SDAP2 | B7FRC2 | 127 | |
| **Membrane distal** | **Phaeo ID** | **Length** | **Abundance** |
| Nad4m/ND4 | F1DGM3 | 493 | |
| Nad5m/ND5 | F1DGQ5 | 671 | |
| AGGG * | B7FUA5 | 123 | |
| B22 | B7FV90 | 83 | |
| SDAP2 | B7FRC2 | 127 | |
| ASHI * | B7GCP1 | 127 | |
| ESSS * | B7FXQ6 | 130 | |
| PDSW * | B7FYP7 | 107 | |

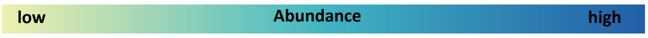


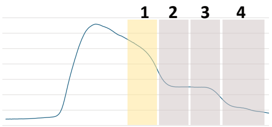


| CII | Phaeo ID | Length | Abundance |
|---|---|---|---|
| SDH1 | B5Y5N6 | 638 | |
| SDH2 | B7GA40 | 273 | |
| SDH3 | B5Y4R0 | 167 | |
| SDH4 | B7G228 | 155 | |

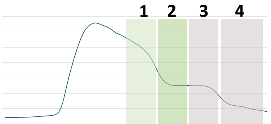


| CIII | Phaeo ID | Length | Abundance |
|---|---|---|---|
| Rieske | B7FYB7 | 179 | |
| CytC1 | B7FVX0 | 272 | |
| QCR6 | B7G063 | 114 | |
| QCR10 * | B7GA54 | 202 | |
| MPP-Beta | B7FYC4 | 115 | |
| QCR7 | B7G150 | 473 | |

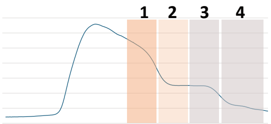


| CIV | Phaeo ID | Length | Abundance |
|---|---|---|---|
| Cox1 | F1DGN0 | 494 | |
| Cox2 | F1DGM7 | 268 | |
| Cox3 | F1DGM5 | 272 | |
| Cox5c | B7G0C7 | 255 | |
| Cox7c | B7FUS0 | 91 | |
| Cox7a * | B7GBE6 | 138 | |

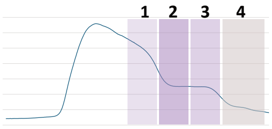


| CV | Phaeo ID | Length | Abundance |
|---|---|---|---|
| ATP beta | B7FS46 | 501 | |
| ATP gamma | B7FTE2 | 305 | |
| ATP epsilon | B7G8W9 | 130 | |
| ATP b | B5Y4I1 | 373 | |
| OSCP | B7FUM8 | 220 | |
| ATP d | B7G9C2 | 688 | |
| ATP 6.8PL * | B7G7I0 | 114 | |
| ATP a | F1DGN2 | 254 | |

**Supplementary Figure 10:** Identification of OXPHOS complexes using sucrose gradient fractionation of enriched isolated mitochondria followed by proteomic analysis

**a,** Summary of membrane enrichment and complexes separation. **b,** Mass-spectrometry (MS) profiling of proteins constituting different OXPHOS complexes across sucrose gradient fractions used to separate isolated mitochondrial complexes. Asterix (*) indicates subunits found exclusively by MS that are unidentifiable by homology search. The 260 nm absorption spectrum used to guide the fractionation is shown, with the fractions indicated and color-coded by the abundance of OXPHOS components.

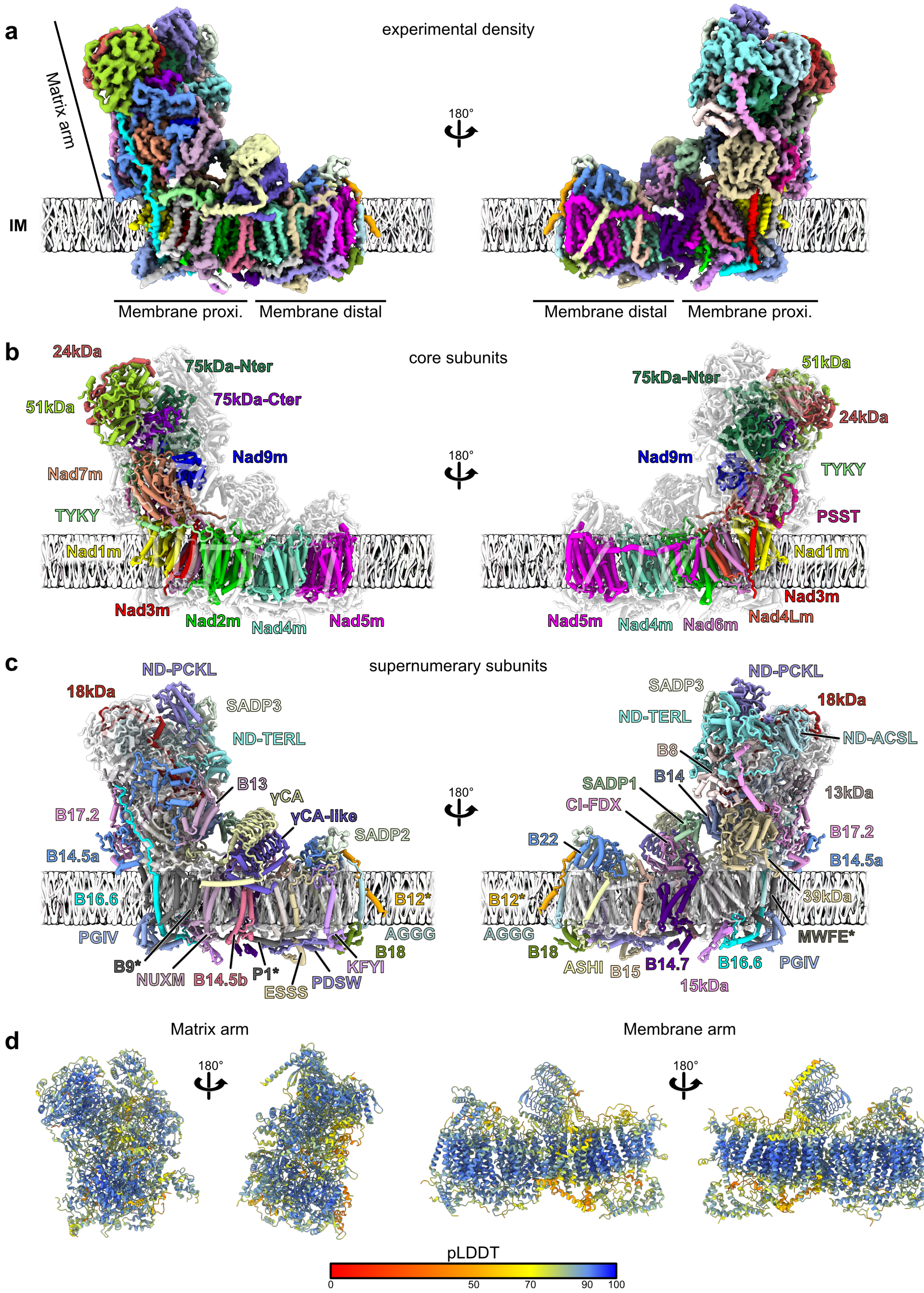


**Supplementary Figure 11:** Model building and composition for complex I

**a,** Subtomogram average map of Phaeodactylum complex I. Each of the 51 proteins are depicted with a different color. The resulting model is shown in **b** and **c**, with **b** depicting the 15 core subunits (with the 75kDa split in two parts) and **c** the annotated supernumerary subunits (36 total). IM stands for inner membrane. The subunits marked with a *, B9, P1, MWFE and B12, were modeled as poly-alanine chains as no annotated candidate could be found in the proteome of either *P. tricornutum* or *T. pseudonana.* **d,** Initial Alphafold3 models of the matrix and membrane arms (separately predicted) depicted with pLDDT values.

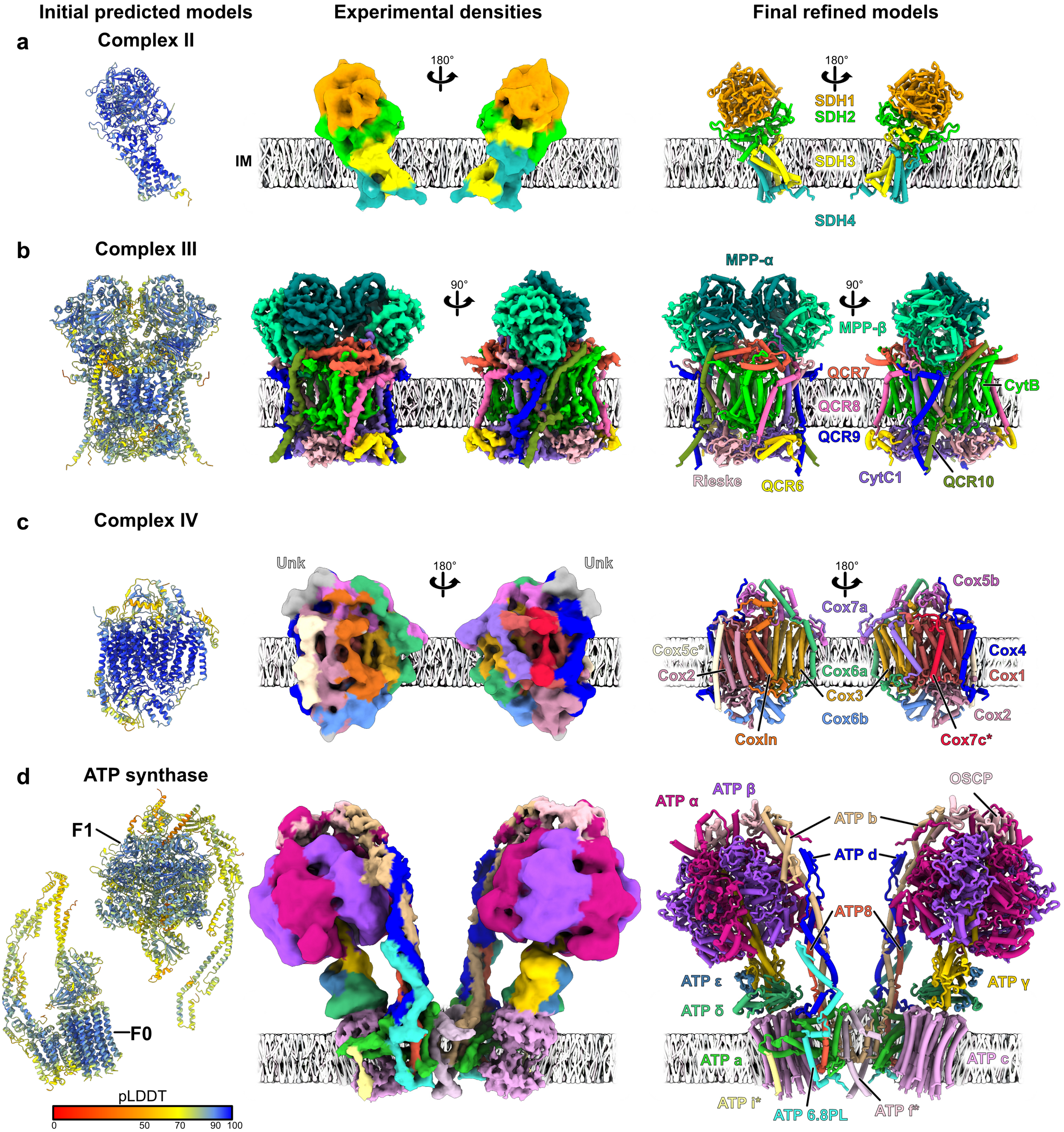


**Supplementary Figure 12:** Model building and composition for respiratory chain complexes II, III, IV and ATP synthase

Overview of the structural determination workflow for Phaeodactylum complex II (**a**), complex III (**b**), complex IV (**c**), and ATP synthase (**d**). Initial AlphaFold predicted models (left) are colored by pLDDT confidence values (scale at bottom). Experimental subtomogram average density maps (middle) and final refined atomic models (right) illustrate the subunit organization within the inner membrane (IM), with all subunits annotated. **a,** Complex II is composed of 4 subunits. **b,** Complex III is a homodimer of 10 subunits. **c,** Complex IV is composed of 11 subunits. The subunits marked with a *****, Cox5c and Cox7c, could not be found in diatom proteomes and were predicted and modeled as yeast subunits. **d,** ATP synthase is a homodimer of 14 subunits and was predicted in two parts. The subunits marked with a *****, ATP i and f, were modeled as poly-alanine chains as no annotated candidate could be found diatom proteomes. i is annotated based on the *Y. lipolytica* structure[57] and f based on Supplementary Fig. 13.

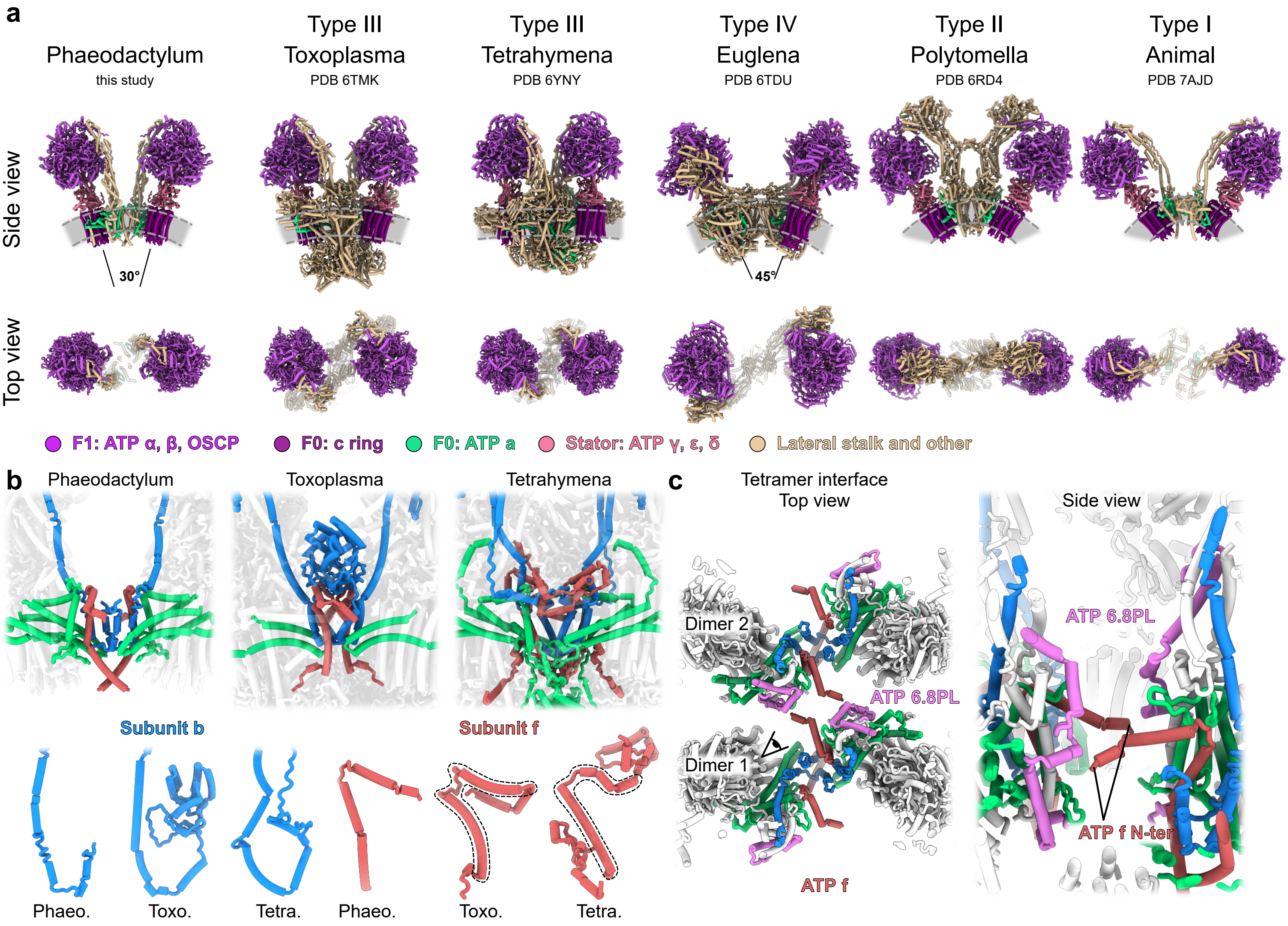


**Supplementary Figure 13:** Structural comparison and dimer interface of the *Phaeodactylum tricornutum* mitochondrial ATP synthase

**a,** Structural comparison of Phaeodactylum ATP synthase against type I, II, III and IV ATP synthases. Phaeodactylum closely resembles Type III architectures (such as those of Alveolates) in overall dimer orientation, and lateral stalk positioning. In terms of subunit composition, however, it remains streamlined, resembling the low subunit complexity of Type I animal complexes, and the inter-monomer angle is of ~30° against 0° in Type III or 45° in Euglena. **b**, Zoomed-in view of the dimer interface comparing Phaeodactylum with Type III ATP synthases. Subunits ATP b (blue) and ATP f (red) are highlighted, with ATP a (green) shown as a structural reference and remaining subunits shown as transparent white density. Despite the acquisition of multiple lineage-specific supernumery subunits in Toxoplasma and Tetrahymena, the conserved core of the dimer interface is established by ATP f and the N-terminal region of ATP b, which contributes to both the lateral stalk and dimer contacts. Isolated atomic models of subunits b and f across the three organisms are shown below for comparison. **c,** Zoomed-in views of the Phaeodactylum tetramer interface, both in top and side views (as seen from the eye pictogram). This highlights that the N-ter of the ATP f subunit extends and mediates some interaction with the ATP 6.8PL subunit.

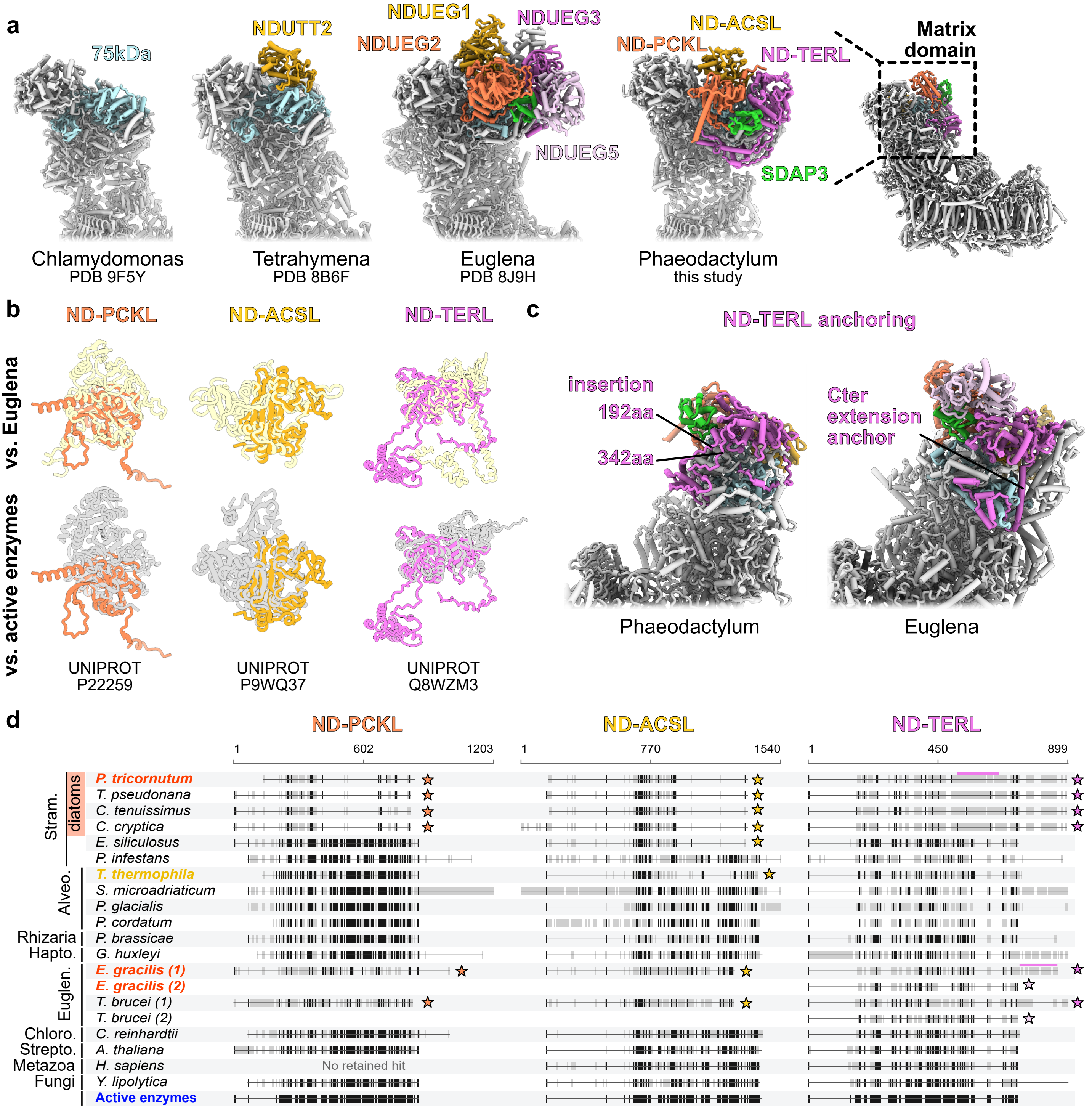


**Supplementary Figure 14:** Structural diversity and evolution of the complex I accessory domain

**a,** Structural comparison of the complex I matrix domain from Phaeodactylum with corresponding domains from Chlamydomonas, Tetrahymena and Euglena. With ND as in NADH Dehydrogenase and PCKL for phosphoenolpyruvate carboxykinase-like, ACSL acyl-CoA synthetase-like and TERL trans-2-enoyl-CoA reductase-like. *C. reinhardtii* lacks additional accessory matrix subunits, resembling the canonical metazoan architecture. *T. thermophila* possesses an ND-ACSL homolog (NDUTT2), whereas *E. gracilis* contains homologs for ND-PCKL (NDUEG2), SDAP3, ND-ACSL (NDUEG1), and an ND-TERL homolog (NDUEG3), plus NDUEG5, which is also a homolog of ND-TERL. The conserved 75 kDa subunit (light blue) is highlighted for structural orientation. **b,** Structural comparison of the Phaeodactylum subunits with their respective Complex I homologs in Euglena (top row) and active enzymes (bottom row). The Phaeodactylum proteins are further reduced compared to Euglena. **c,** Detailed view of the differentially anchored ND-TERL subunits, illustrating divergent structural adaptations for matrix arm attachment via an internal anchor insertion in Phaeodactylum versus a C-ter extension anchor in Euglena. **d,** Sequence alignments of the identified homologous subunits across major eukaryotic lineages (protein IDs listed in Supplementary Table 3) compared with active reference enzymes (*E. coli* PDB 1AQ2, *M. tuberculosis* PDB 3R44, *C. tropicalis* PDB 4WAS). Organisms where the subunits were identified are highlighted in yellow and orange. Grey blocks represent residues, with black ticks indicating sequence identity to the active enzymes. *C. reinhardtii, A. thaliana, H. sapiens* and *Y. lipolytica* do not have these additional subunit and act as negative controls. Established complex I-associated homologs are marked with a star. The three accessory subunits are conserved across all analyzed diatoms. Among other stramenopiles, *E. siliculosus* retains only the ND-ACSL module, similar to *T. thermophila* in Alveolata, while the heterotrophic oomycete *P. infestans* appears to lack these subunits entirely. For ND-TERL, extensions involved in anchoring are highlighted.

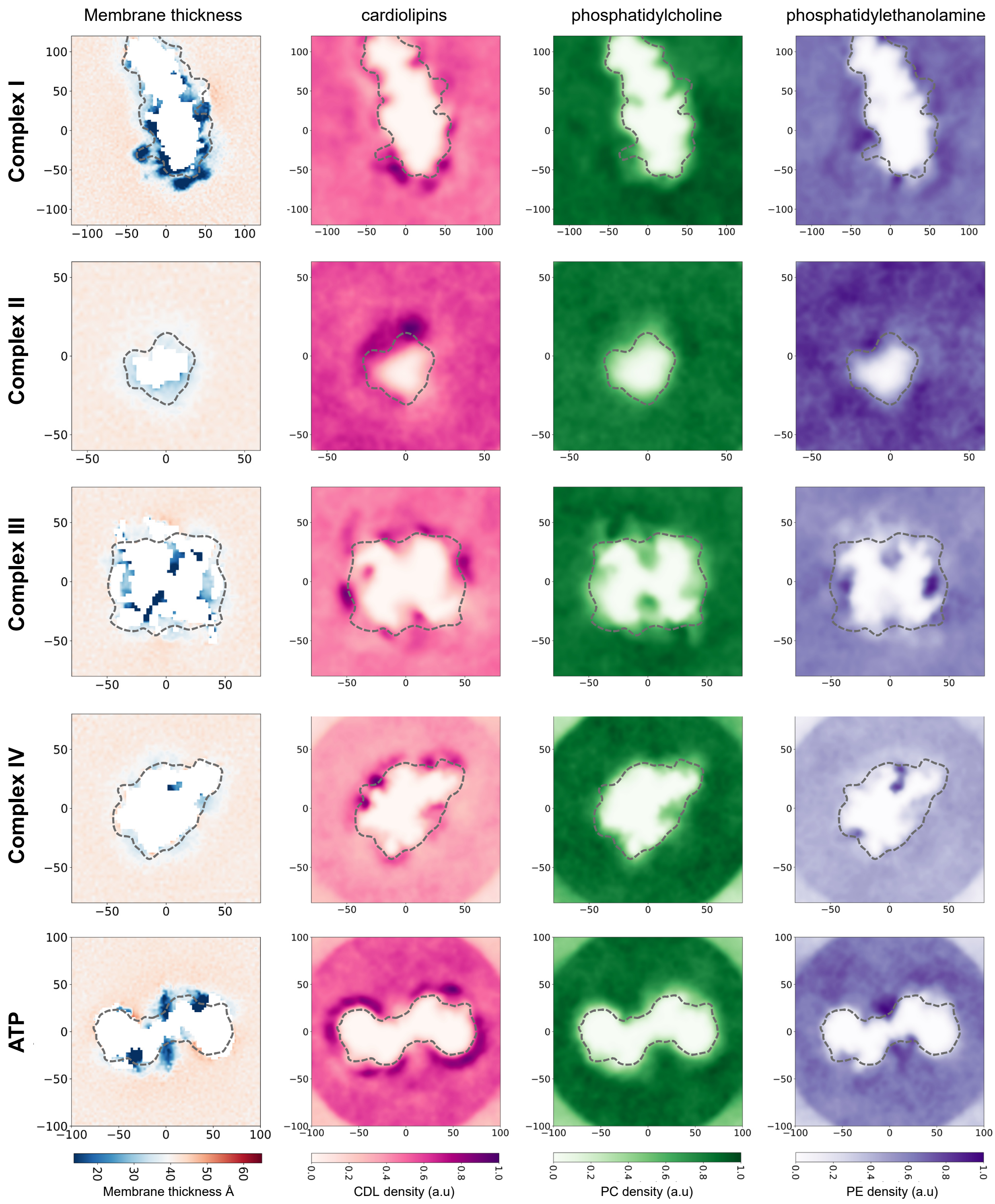


**Supplementary Figure 15:** Membrane thickness and lipid density distributions around respiratory chain complexes and ATP synthase from coarse-grained molecular dynamics simulations

Each row displays results for a specific mitochondrial complex: Complex I, Complex II, Complex III, Complex IV, and ATP synthase. Columns (left to right): Membrane thickness, calculated in Å. Cardiolipin (CDL) density are presented in arbitrary units (a.u.) from 0.0 to 1.0 in pink to purple. Similarly, phosphatidylcholine (PC) densities are presented from light to dark green and phosphatidylethanolamine (PE) densities are shown in light to dark purple. Strong membrane thinning is observed around complex I and all complexes preferentially recruit negatively charged cardiolipin. Systems were modeled using the Martini 3 force field and embedded in a symmetric PE:PC:CDL (4:2:1) lipid bilayer. For each complex, five independent 5 µs production simulations were performed. The dashed grey contour in each panel outlines the perimeter of the respective protein complex.
